# U-rich elements drive pervasive cryptic splicing in 3’ UTR massively parallel reporter assays

**DOI:** 10.1101/2024.08.05.606557

**Authors:** Khoa Dao, Courtney F. Jungers, Sergej Djuranovic, Anthony M. Mustoe

## Abstract

Non-coding RNA sequences play essential roles in orchestrating gene expression. However, the sequence codes and mechanisms underpinning post-transcriptional regulation remain incompletely understood. Here, we revisit the finding from a prior massively parallel reporter assay (MPRA) that AU-rich (U-rich) elements in 3’ untranslated regions (3’ UTRs) can drive upregulation or downregulation of mRNA expression depending on 3’ UTR context. We unexpectedly discover that this variable regulation arises from widespread cryptic splicing, predominately from an unannotated splice donor in the coding sequence of GFP to diverse acceptor sites in reporter 3’ UTRs. Splicing is activated by U-rich sequences, which function as potent position-dependent regulators of 5’ and 3’ splice site choice and overall splicing efficiency. Splicing has diverse impacts on reporter expression, causing both increases and decreases in reporter expression via multiple mechanisms. We further provide evidence that cryptic splicing impacts between 10 to 50% of measurements made by other published 3’ UTR MPRAs. Overall, our work emphasizes U-rich sequences as principal drivers of splicing and provides strategies to minimize cryptic splicing artifacts in reporter assays.

## INTRODUCTION

Post-transcriptional regulation of RNA splicing, polyadenylation, translation, and degradation plays essential roles in shaping gene expression^1^. These post-transcriptional regulatory programs are predominantly encoded by non-coding RNA sequences located in intronic and untranslated regions (UTRs) that recruit diverse trans-acting factors, including small nuclear RNAs, RNA-binding proteins (RBPs) and microRNAs (miRNAs)^2,3^. Dysfunction of splicing and other post-transcriptional regulatory processes has been implicated in diverse human diseases^1^. However, the molecular codes that prescribe splicing and post-transcriptional regulation remain cryptic^4,5^. Understanding the functions and mechanisms of non-coding sequences continues to be a critical goal in biology.

Reporter assays have long been one of the most important strategies for studying post-transcriptional regulation. These assays place a reporter gene such as GFP under the control of non-coding regulatory sequences^6^, enabling isolated measurement of non-coding sequence function. Recently, reporter assays have been extended to permit functional evaluation of thousands of non-coding sequences in parallel, termed massively parallel reporter assays (MPRAs)^6^. MPRAs leverage advances in gene-synthesis technology to clone large libraries of non-coding sequences into a common reporter vector in a pooled format. The pooled library is then transfected into cells and the activity of each non-coding sequence is measured via targeted next-generation sequencing. The flexibility and high throughput of MPRAs make them useful for addressing diverse questions, including annotating the function of non-coding regulatory elements, evaluating the impact of genetic variants, and comparing homologous regulatory elements across species^7–13^. However, recent studies have identified multiple, frequently overlooked design choices that can convolute MPRA measurements^14^. For instance, the same candidate regulatory element can exhibit significantly different activity depending on whether the element is cloned with more or less of its endogenous surrounding sequence context^14^. Transfecting MPRA libraries as episomal vectors versus integrating them into chromatin using viral vectors, or use of alternative sequencing strategies to measure MPRA expression, can also generate different answers^15^. Fully defining the limitations and design caveats of MPRAs is important for ensuring that these experiments yield faithful measurements of non-coding sequence function.

Cryptic splicing has long been known to a be an artifact that can plague reporter assays, causing alteration of reporter expression, false-positive identification of internal ribosome entry sites (IRESes), and production of fusion proteins, among other examples^16–19^. Cryptic splicing arises when an inserted sequence element activates splicing from unexpected, usually weak 5’ donor or 3’ acceptor sequences. However, the requirements for activating cryptic splice sites remain poorly understood and are challenging to predict^20–22^. MPRAs may be particularly susceptible to cryptic splicing due to architectural features of reporter plasmids and the diversity of sequences that are assayed. Additionally, it is common for MPRA experiments to only sequence short segments of each reporter, meaning that cryptic splicing may go undetected. Better understanding the mechanisms underpinning cryptic splicing can help improve MPRA designs. Furthermore, studying cryptic splicing in the MPRA context may potentially offer insights into how mis-splicing arises in human disease contexts^23^.

One of our groups recently used an MPRA-based strategy to evaluate combinatorial regulatory interactions between 3’ UTR sequence motifs^24^. This strategy, termed post-transcriptional regulatory element sequencing (PTRE-seq), employed a library of synthetic 3’ UTRs that each of encoded an array of four regulatory modules. Each module coded for either a “blank” control sequence, a let-7 miRNA site, Pumilio protein recognition element (PRE), Smaug protein recognition element (SRE), or an AU-rich element (ARE), which are bound by diverse ARE-binding RBPs. All combinations of these regulatory modules were synthesized and cloned into the 3’ UTR of a common GFP reporter, resulting in a library of 642 unique reporters. PTRE-seq revealed that these regulatory elements generally function additively to repress mRNA stability and translation. However, as a surprising exception, AREs exhibited strong epistatic interactions with both adjacent AREs and other regulatory motifs, sometimes dramatically enhancing or repressing reporter expression depending on their relative position in the 3’ UTR. The mechanism for this variable, position-dependent impact of AREs on gene expression was unclear, but pointed to potential contextual control of 3’ UTR regulation^25^.

In this work, we revisited the PTRE-seq MPRA experiment to better define the mechanism for position-dependent regulation of AREs. Unexpectedly, we discovered that PTRE-seq reporters undergo widespread cryptic splicing invisible to conventional MPRA measurement strategies. Cryptic splicing utilizes common GFP sequences and is activated by 3’ UTR ARE (U-rich) elements in a position-dependent manner, explaining the variable ARE phenotype. Repurposing this PTRE-seq dataset revealed novel aspects of splicing regulation, including insights into how U-rich elements modulate splice site selection and splicing efficiency, as well as the impact of splicing on transcription. Additionally, we show that cryptic splicing impacts conclusions drawn by other MPRAs of 3’ UTR function. Overall, our work emphasizes the central role of U-rich elements in driving splicing and provides preventative and corrective solutions for minimizing splicing artifacts in MPRA designs.

## RESULTS

### The PTRE-seq MPRA library features widespread cryptic splicing

As part of an effort to understand position-dependent ARE regulation, we repeated the PTRE-seq experiment^24^ with slight modifications. HeLa cells were transfected with the same PTRE-seq plasmid library followed by total RNA isolation and amplicon sequencing to quantify reporter abundance. In the original experiment, reporter abundance was quantified via targeted sequencing of short co-transcribed barcode elements, which is a standard strategy used in MPRA experiments (“barcode primers”, Fig. 1A)^6^. By contrast, we used a pair of “extended” primers to obtain sequencing coverage across most of the 3’ UTR and into the GFP CDS (Fig. 1A). Unexpectedly, the amplicon libraries obtained using these extended primers exhibited numerous shorter products, contrasting with the uniform amplicon size obtained when preparing libraries from the parent DNA plasmids (Fig. 1B). RNA abundance measurements obtained using extended primers were well-correlated with published barcode-based measurements (R = 0.73; Fig. S1A), supporting that our experiments faithfully replicate the original PTRE-seq assay. Thus, these results indicated that mRNA transcripts in the PTRE-seq library have significant, unappreciated size heterogeneity.

**Figure 1.**
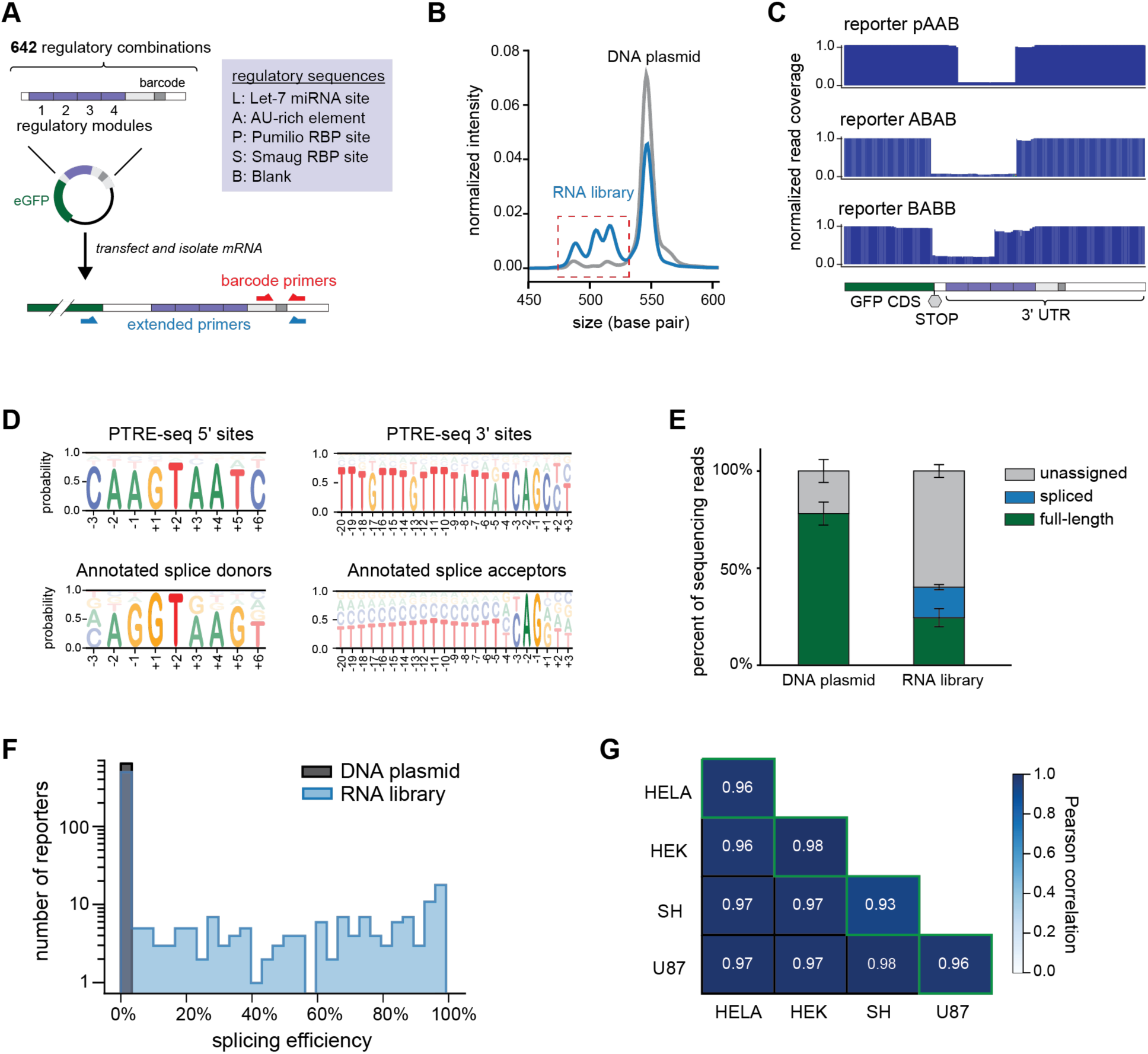
The PTRE-seq MPRA library undergoes pervasive cryptic splicing. (A) Schematic of the PTRE-seq library. Original “barcode” primers and “extended” primers used in this work are shown at bottom. eGFP, enhanced green fluorescent protein. (B) Size heterogeneity was observed in sequencing libraries prepared from RNA (blue) but not DNA plasmid library (grey). Electropherogram was measured by Tapestation 4200. (C) Representative read coverage tracts of reporters with internal deletions. (D) Position weight matrices of sequences observed at 5’ and 3’ deletion boundaries observed in PTRE-seq reporters and annotated 5’ and 3’ splice sites retrieved from the human GRCh38 genome assembly. (E) Percent of PTRE-seq reads that are full-length, spliced, or unassigned from DNA and RNA sequencing libraries. Error bars denote standard deviation across two biological replicates. (F) Distribution of splicing efficiencies of PTRE-seq reporters. (G) Pearson correlations of PTRE-seq splicing efficiencies measured across biological replicates of the same cell line (diagonal, green outline) and between different cell lines.

Although PTRE-seq reporters were not designed to be spliced and lack any known splice sites, sequencing of the extended PTRE-seq RNA library strongly suggested that this size heterogeneity was due to cryptic splicing within reporter 3’ UTRs. A subset of reporters featured internal deletions with precisely defined boundaries (Fig. 1C). Sequence analysis of the 5’ and 3’ deletion sites revealed strong enrichment for sequences that resemble splice sites, including characteristic GT and AG dinucleotides at position +1 of the 5’ site and -2 of the 3’ site respectively (Fig. 1D)^4^. U-rich regions were also strongly enriched upstream of the 3’ site, consistent with a polypyrimidine tract^26^. We validated these MPRA-based observations with RT-PCR analysis on selected individually transfected reporters, which confirmed the presence of novel splice junctions in the 3’ UTR of PTRE-seq transcripts (Fig. S1B).

To quantify the frequency of splicing and other potential artifacts in our MPRA sequencing library, we developed a computational pipeline to categorize PTRE-seq reads either as “full length”, “spliced”, or “unassigned” (Fig. S1C). Full length reads contain concordant UTR sequences and barcodes that match a known reporter design. Spliced reads contain concordant UTR sequences and barcodes but also feature an extended internal deletion bounded by +1 GT and -1 AG dinucleotides. The remainder of reads were considered “unassigned” and had either internal deletions not bounded by GT and AG, unassignable barcodes (possibly due to being partially or completely spliced out), or had discordant UTR and barcode combinations suggestive of recombination during reverse-transcription or PCR (Fig. S1D, S1E). Usage of an alternative reverse-transcriptase during library preparation resulted in fewer apparent recombination artifacts without significantly impacting splicing measurements (MarathonRT^27^ compared to Superscript II; Fig. S1F, S1G). Overall, 41% of RNA library reads were assignable, compared to 68% for the DNA plasmid library (Fig. 1E). These analyses emphasize the need for thoughtful quality control during library preparation and bioinformatic quantification^14^.

Quantification of the fraction of spliced versus full-length reads observed for each reporter revealed significant reporter-specific cryptic splicing (Fig. 1F). Out of 642 reporters, 119 (18%) were spliced at 20% or higher rates and 32 (5%) were nearly completely spliced (> 90%). Reporter splicing efficiencies were reproducible between biological replicates (Pearson correlation = 0.98) (Fig. 1G, S1H) and across internal barcode replicates (Fig. S1I). Splicing efficiencies were also in agreement with orthogonal quantification by semi-quantitative RT-PCR of individually transfected reporters (Fig. S1B). We also explored whether this cryptic splicing varied across different cell types. Splicing efficiencies were highly consistent across experiments performed in three additional cell lines: human embryonic kidney (HEK293), human neuroblastoma (SH-SY5Y), and human glioblastoma (U87) (Fig. 1G). Thus, cryptic splicing in the PTRE-seq library is constitutive and generalizable across diverse cell contexts.

### Splicing occurs at diverse weak donor and acceptor sites

Given that the PTRE-seq library was not designed to contain splice sites, we sought to better understand the sequence features driving efficient 3’ UTR splicing. Strikingly, 127 out of 150 spliced reporters (>1% splicing efficiency) use a cryptic 5’ splice site located at the stop codon of the eGFP coding sequence (Fig. 2A). Splicing also occurred from a donor located two codons upstream of the stop codon, albeit at significantly reduced efficiency (alt-GFP; Fig. 2A). Splicing at these 5’ sites excises the stop codon, resulting in a C-terminal extension of GFP protein that was detectable by Western Blot (Fig. S2A). These C-terminal-extended GFP proteins exhibit significantly lower expression than expected based on RNA abundance, consistent with the known destabilizing effect of C-terminal protein extensions (Fig. S2B)^28^. While both the GFP and alt-GFP donor sites contain GT dinucleotides, they are otherwise weak sites compared to known human splice donors (1^st^ percentile for both donors; Fig. 2B). Nevertheless, the GFP donor was able to support nearly 100% splicing efficiency of multiple reporters (Fig. S2C).

**Figure 2.**
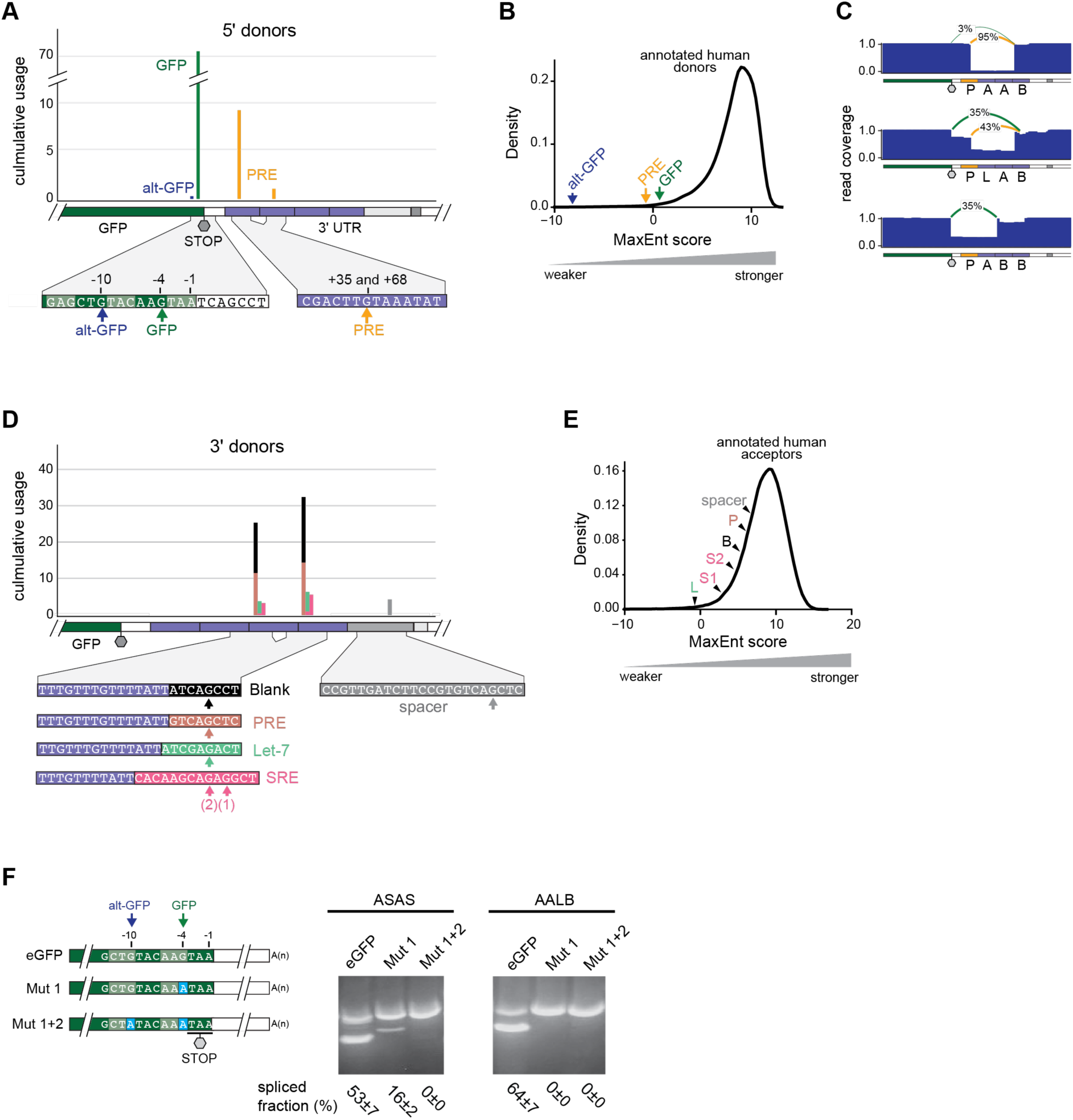
Diverse weak splice donors and acceptors are used in the PTRE-seq library. (A) Location and usage frequency of splice donors in PTRE-seq reporters. Cumulative usage was computed as the sum of splicing efficiencies for each splice donor across all reporters. Nucleotide sequences are detailed in the diagram below. (B) 5’ donor site strengths of PTRE-seq splice donors and annotated human splice donors. (C) Representative read coverage tracts illustrating alternative usage of the GFP and PRE donors. (D) Location and usage frequency of splice acceptors in PTRE-seq reporters. Cumulative usage was computed as the sum of splicing efficiencies at each corresponding splice acceptor across all reporters. Sequence details of each acceptor are shown at bottom. (E) 3’ acceptor site strengths of PTRE-seq splice acceptors and annotated human splice acceptors. (F) Synonymous mutations that abolish GFP cryptic splice donors abrogate splicing in representative reporters. Schematics of mutations are shown at left. RT-PCR analysis of recoded ASAS and AALB reporters is shown at right. Reporters were individually transfected into HeLa cells, followed by RT-PCR and resolved on an agarose gel. Quantification shown below the gel represents the median and standard deviation over 3 biological replicates. For (B) and (E), splice site strength is quantified by using Maximum Entropy score ^52^.

In addition to the GFP coding sequence donors, we also observed usage of a second set of cryptic 5’ splice sites in the 3’ UTRs of 23 reporters. These sites mapped to PRE regulatory modules present in either the 1^st^ or 2^nd^ positions of the 3’ UTR regulatory array (Fig. 2A, 2C). Like the GFP sites, these donors are weak (1^st^ percentile relative to annotated human splice donors; Fig. 2B) but were capable of supporting >90% efficiency depending on neighboring sequence context (Fig. S2C). Identical P modules located in the 3^rd^ or 4^th^ module positions were not used as donors (Fig. S2D), which is likely explained by a lack of appropriately spaced downstream acceptor sites. Notably, because all PTRE-seq reporters contain potential GFP CDS donors, usage of 3’ UTR P donors occurs in competition with GFP, and complex patterns of alternative splicing were observed in some P-containing reporters (Fig. 2C, S2D).

Given that all PTRE-seq transcripts contain viable 5’ splice sites (the GFP CDS), we hypothesized that cryptic splicing might be determined by the availability and strength of downstream 3’ acceptor sites. However, 3’ splice acceptors exhibited minimal sequence dependence. 3’ splice sites predominantly mapped to the 3^rd^ and 4^th^ regulatory modules with all regulatory elements except for AREs serving as splice acceptors (Fig. 2D). We also observed a subset of reporters that utilized a 3’ acceptor site located downstream of the regulatory array, in one of the constant spacer regions (Fig. 2D). All 3’ splice sites featured an AG dinucleotide and an upstream U-rich region (Fig. 2D). By contrast, ARE modules lack an AG dinucleotide motif, explaining why they are not used as acceptors. Similar to 5’ donors, these 3’ splice acceptors are substantially weaker than annotated human 3’ donors (1^st^ to 28^th^ percentile, Fig. 2E). Acceptor site strength also poorly correlated with splicing efficiency (Fig. S2E). The restriction of splicing to the 3^rd^ and 4^th^ regulatory modules but not the 1^st^ and 2^nd^ module positions can be explained by the 70-nucleotide minimum intron size needed for spliceosome assembly (Fig. S3F)^29^. Altogether, our data indicate that factors beyond 5’ and 3’ splice site availability govern splicing in the PTRE-seq library.

To validate the function of the identified GFP splice sites, we used site-directed mutagenesis to synonymously recode the GFP C-terminus in several highly spliced reporters. Removal of the primary GFP donor via an AAG→AAA substitution at the -2 codon suppressed most splicing, but residual splicing was still observed at the secondary alt-GFP site (Fig. 2F). Full suppression was achieved with an additional CTG→CTA substitution at the -4 codon. Notably, these recoded GFP proteins were expressed at equivalent or greater levels than standard GFP (Fig. S2G), indicating that these splice-suppressing mutations do not significantly impact translation. Overall, these results emphasize that common sequences can serve as cryptic splice sites and drive complex splicing behaviors.

### AU-rich elements activate cryptic splicing and determine splice site choice in the PTRE-seq library

Our analyses indicate that every reporter contains both feasible 5’ and 3’ splice sites, yet splicing efficiencies varied dramatically across reporters. Further analysis revealed that splicing was almost entirely determined by the presence and locations of ARE modules. Splicing strictly depends on the presence of at least one 3’ UTR ARE module within the cryptic intron (Fig. 3A, B). For some splice acceptor sites, such as B and P, a single ARE module is sufficient to activate strong splicing (Fig. 3C, S3A). For example, reporter BBBB is unspliced, but reporters BABB and BBAB are spliced with >80% efficiency (Fig. S3B). An increased number of intronic ARE modules also drives increased splicing efficiency at all acceptors, including activating weaker splice acceptors (Fig. 3C, S3A). For instance, L, S, and spacer acceptors are only used when there are at least two upstream AREs (Fig. 3C, S3A). This ability of multiple AREs to increase splicing efficiency was observed regardless of whether AREs were contiguous or non-contiguous (Fig. S3C).

**Figure 3.**
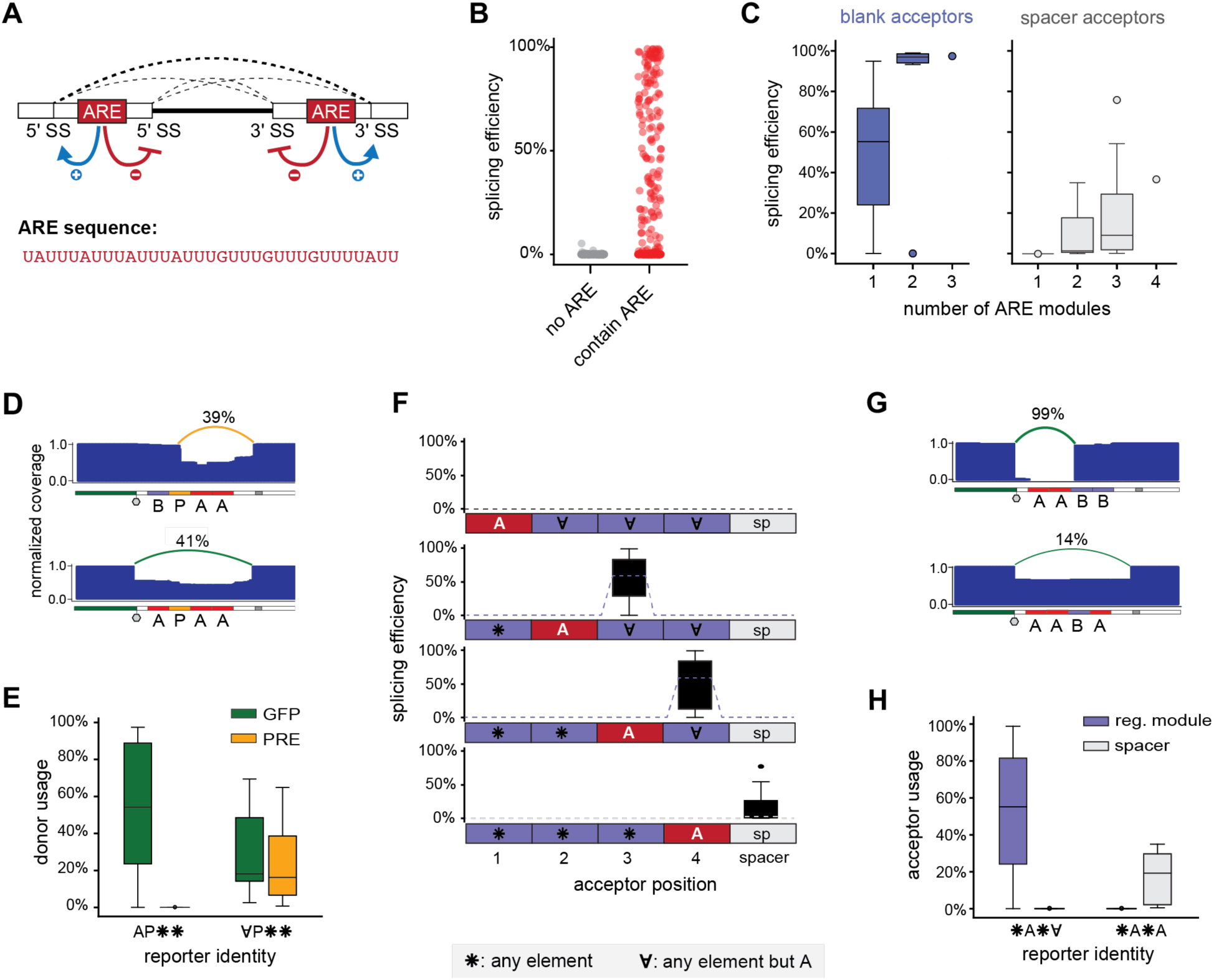
AU-rich elements (AREs) are potent splicing regulators in the PTRE-seq library. (A) The nucleotide sequence of the PTRE-seq ARE element and a cartoon illustrating its impact on splice site selection. (B) Splicing efficiencies of ARE-containing and non-ARE containing PTRE-seq transcripts. (C) Relationship between the number of intronic AREs and splicing efficiency for blank and spacer splice acceptors. (D) Example read coverage tracts illustrating ability of 5’ AREs to block usage of adjacent PRE donors. (E) Comparison of donor usage in reporters with and without AREs 5’ to a 2^nd^ position PRE donor. (F) Splice acceptor usage based on location of the 3’-most ARE module. Box plots represent the distribution of splicing efficiencies observed at each potential acceptor site (1^st^, 2^nd^, 3^rd^, or 4^th^ module sites, or spacer site) for different combinations of regulatory elements denoted along the bottom. (G) Example read coverage tracts illustrating ability of 3’ AREs to block usage of a strong upstream splice acceptor. (H) Comparison of donor usage in reporters with and without AREs in the 4^th^ regulatory module. For (C, E, F, H), each box plot represents the distribution of spliced fraction for a group of reporters. Whiskers indicate the furthest datum that is 1.5*Q1 (upper) or 1.5*Q3 (lower). * symbol wrepresents all regulatory elements, 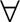 symbol represents all regulatory elements except ARE.

Out data also reveal that AREs play a major role in dictating both 5’ and 3’ splice site choice. At the 5’ donor site, an increased number of “intronic” AREs promotes selection of P donors over GFP donors (Fig. S3D, S3E). For instance, reporters PSAB and PAAB both contain a potential GFP donor and 1^st^ module P donor and share identical 3’ acceptor sites. PSAB is predominantly spliced at the GFP donor, but PAAB is almost exclusively spliced at the P donor (Fig. S3D). However, AREs that are 5’-adjacent to potential donors (i.e. are exonic) are repressive of splice site selection. For example, reporter BPAA is exclusively spliced using a P donor whereas the closely related APAA reporter switches to the GFP donor site (Fig. 3D). This observation generalizes across reporters, with an ARE in the 1^st^ position blocking usage of 2^nd^ position P sites (Fig. 3E). Thus, AREs function as exonic silencers and intronic enhancers of 5’ splice sites.

For the majority of reporters, the 3’ splice site is located immediately downstream of the 3’-most ARE, reflecting that the ARE module likely functions as a polypyrimidine tract (Fig. 3F). When AREs are located in the 2^nd^ or 3^rd^ module, splicing occurs at the most proximal AG dinucleotide in the 3^rd^ or 4^th^ position, respectively (Fig. 3F). Similarly, in reporters where the 3’-most ARE is at the 4^th^ position, the acceptor site shifted to the most proximal AG site in the adjacent spacer sequence (Fig. 3F). As exceptions to this rule, reporters with 3’-most AREs in the 1^st^ position go unspliced, likely because the resulting intron would be less than 70 nucleotides (Fig. S2F). The relative spacing between the ARE polypyrimidine tract and the downstream GA site also explained the difference in splicing efficiency of different acceptor modules^26^: B and P modules features GA sites only 5 nucleotides downstream from an adjacent ARE, whereas AG sites are located 6 and 9 nucleotides downstream in L and S modules (Fig. 2D). For the poorest splicing spacer acceptor, the nearest AG site is 30 nucleotides downstream of a 4^th^ position ARE module. Spacer acceptors thus likely use a shorter and less U-rich polypyrimidine tract encoded within the spacer (Fig. 2D).

Interestingly, the rule that splicing occurs downstream of the 3’-most ARE applied even when superior splicing sites were available upstream (Fig. 3G). For instance, reporter AABB is >98% spliced using the favorable 3^rd^ position B acceptor. By comparison, reporter AABA, which contains the same 3^rd^ position B site, instead switches to the weaker spacer acceptor site and is spliced at only 14% efficiency (Fig. 3H). Thus, AREs can suppress usage of upstream acceptor sites, inducing switching to weaker downstream acceptors. Overall, our data demonstrate that “intronic” AREs are potent activators of cryptic splicing, whereas “exonic” AREs can suppress both 5’ and 3’ splice site selection (Fig. 3A).

### Cryptic splicing modulates reporter expression via multiple mechanisms and explains the position-dependent regulatory effects of AREs

We next explored how cryptic splicing impacts PTRE-seq expression measurements. PTRE-seq was designed to measure changes in gene expression caused by post-transcriptional regulation by 3’ UTRs, but cryptic splicing may modulate expression and consequently convolute result interpretation. Indeed, reporter expression was strongly positively correlated with splicing efficiency (R = 0.74), with the most highly spliced reporters exhibiting a >4× higher expression than baseline “blank” 3’ UTRs (Fig. 4A).

**Figure 4.**
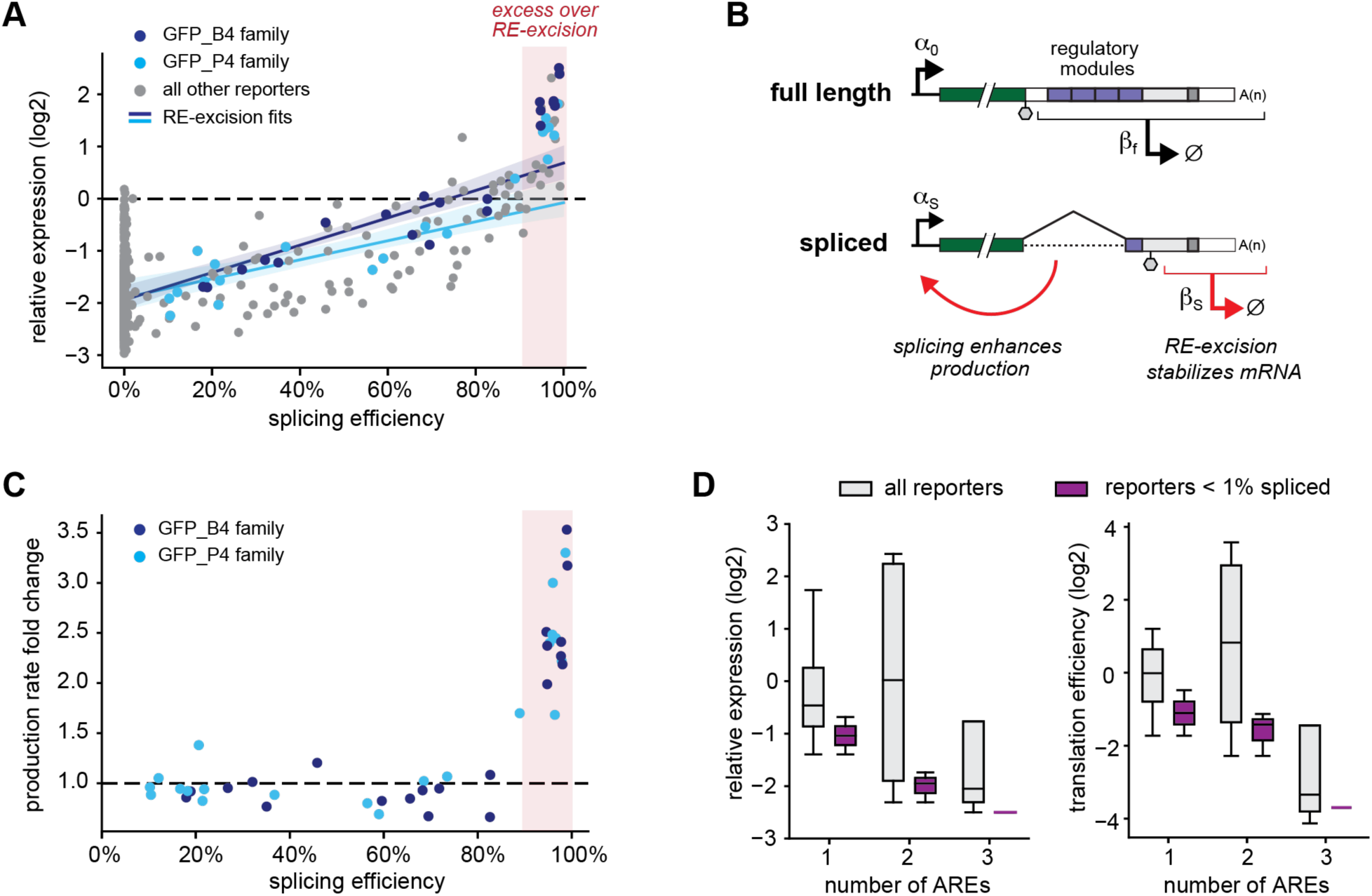
Cryptic splicing enhances PTRE-seq reporter expression via RE-excision and enhanced production. (A) Relationship between splicing efficiency and reporter expression. GFP_B4 and GFP_P4 families of spliced transcripts are shown in dark and light blue, respectively. Regression lines of respective colors show the fits of RE-excision model to GFP_B4 and GFP_P4 reporters. The RE-excision model assumes constant production and is computed via Equation 2 (Methods) based on estimated stabilities of full-length and spliced isoforms (*β_f_* and *β_s_*, respectively). *β_s_* was estimated from Equation 3 using reporters spliced with 10% to 80% efficiencies. Expression measurements are from the original PTRE-seq study^24^ using barcode primers and normalized to the BBBB reporter. (B) Illustration of the mechanisms through which splicing impacts PTRE-seq reporter expression. (C) Estimated production rate for GFP_B4 and GFP_P4 reporters as a function of splicing efficiency. Production rate is computed via Equation 4 (Methods) based on total expression, splicing efficiency, and estimated stabilities of full-length and spliced isoforms (*β_f_* and *β_s_*, respectively). *β_s_* was estimated as in (A). Alternative estimates of *β_s_* gave qualitatively similar results (Fig. S4). (D) Effect of ARE module copy number on RNA steady state expression and translation efficiency. Shown are reporters consisting of all arrangements of blank modules with the indicated number of ARE modules. The original PTRE-seq study was unaware of cryptic splicing, making it appear that AREs have widely divergent impacts on mRNA stability and translation (grey boxplots). When spliced reporters are excluded, AREs have a uniformly destabilizing impact on reporter expression and translation efficiency (purple boxplots). Boxplot whiskers indicate the furthest datum that is 1.5*Q1 (upper) or 1.5*Q3 (lower).

One mechanism through which splicing may increase reporter expression is by excising otherwise destabilizing 3’ UTR regulatory elements (Fig. 4B). We developed a simple “RE-excision” model that describes reporter expression as a sum of full-length and spliced isoforms that have differing stabilities. To fit this model, we focused on groups of reporters that are spliced at identical 5’ and 3’ sites, yielding identical spliced isoforms but with varied splicing efficiency because of different cryptic intronic sequence elements (Fig. S4A). For example, 21 reporters are spliced from a GFP donor to a 4^th^ position blank sequence (GFP_B4), with splicing efficiencies ranging from 18% to 99% (Fig 4A, Fig. S4A). The stability of the full-length isoform of each reporter can be estimated from its combination of 3’ UTR regulatory elements (Methods). The stability of the common spliced isoform (GFP_B4) can then be estimated by regressing the observed total expression of each reporter versus its splicing efficiency (Methods). Fitting this model to two different families of reporters indicated that spliced 3’ UTR isoforms are 10-70% more stable than baseline “blank” 3’ UTRs (Fig. S4B-D). Accounting for RE-excision recapitulated the expression pattern observed for reporters spliced at <90% efficiency (Fig. 4A).

While the RE-excision model explains the expression of reporters spliced at intermediate efficiency, it is unable to explain the dramatic increase of expression above the “blank” baseline for highly spliced reporters (Fig. 4A). We postulated that this increase reflected intron-mediated enhancement of reporter production (Fig. 4B)^30–33^. We solved for the relative production rate needed to explain the expression of each reporter using the full-length and spliced isoform stabilities obtained from RE-excision modeling (Methods). Low-to-moderately spliced reporters feature a constant production rate matching that of unspliced, reference transcripts (Fig. 4C). By comparison, our analysis indicates that reporters with splicing efficiencies >90% exhibit a 2 to 4-fold enhancement in production rate (Fig. 4C). This general trend of production rate increasing non-linearly at splicing efficiencies >90% was consistent across two independent families of reporters (Fig. 4C) and was robust to alternative model fitting strategies (Supplementary Fig. S4E, S4F). Thus, cryptic splicing enhances reporter expression both via relieving RE-mediated destabilization and enhancing the production rate of highly spliced reporters.

Given the observation that ARE-directed splicing can significantly enhance reporter expression, we revisited our prior conclusion that AREs can sometimes stabilize mRNAs and enhance translational efficiency^24^. Limiting analysis to unspliced 3’ UTRs revealed that AREs induce a uniform reduction in mRNA expression and translation efficiency that scales with the number of ARE motifs (Fig. 4D). This observation aligns with the canonical role of AREs as destabilizing elements^34,35^ and indicates that cryptic splicing likely explains the variable ARE effects observed in our original study. We also re-examined our prior conclusion that AREs can either enhance or antagonize the activity of adjacent PRE and let-7 regulatory modules depending on their relative positioning^24^. Regression modeling revealed that these variable epistatic interactions are also artifacts of splicing, with AREs weakly antagonizing PRE and let-7 elements regardless of positioning in unspliced reporters (Fig S5A, S5B). Other conclusions from the PTRE-seq study, which focused on the independent effects of PRE, let-7, and SRE sites, are not impacted by splicing^24^.

### Cryptic splicing artifacts are common in other published MPRAs

Given that splicing in the PTRE-seq library only requires a GFP coding sequence and an ARE (U-rich) sequence, we investigated whether similar splicing artifacts are present in other published MPRAs. To do so, we developed a strategy to predict cryptic splicing using SpliceAI, a deep learning-based tool for predicting 5’ and 3’ splice sites in the human genome (Fig. S6A)^36^. SpliceAI accurately predicted the position of the observed 5’ and 3’ splice sites for the large majority of spliced reporters in the PTRE-seq library (Fig. S6B). To predict total splicing probability of each reporter, we computed the product of the maximum 5’ and 3’ SpliceAI site probabilities. This total splicing score strongly correlated with observed splicing efficiency in PTRE-seq (Fig. 5A). A splicing probability <0.1 reliably identifies unspliced reporters, whereas a >0.6 splicing probability provides a specific predictor of efficient splicing. However, intermediate splicing probabilities have only moderate predictive value; for example, at predicted splicing probability of 0.3, reporters are equally likely to be highly spliced (>90%) as they are unspliced (<1%). The reduced predictive power of intermediate splicing probabilities is consistent with previous evaluations of SpliceAI^37^. We thus used total predicted splicing probabilities of >0.3, >0.6, and >0.9 as predictions of moderate, strong, and very strong splicing, respectively (Fig. 5A).

**Figure 5.**
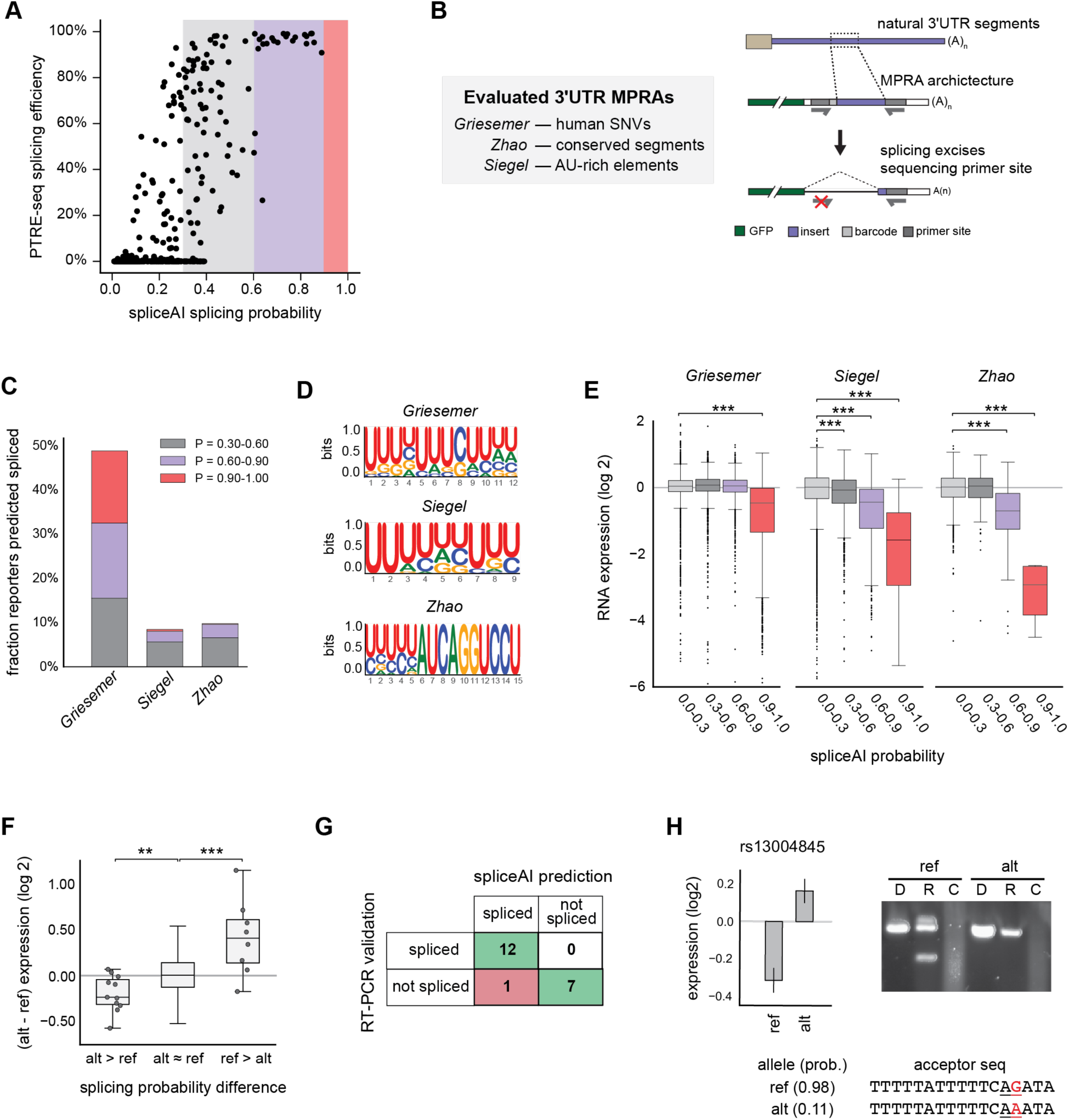
Cryptic splicing is common in published MPRAs. (A) SpliceAI^36^ total predicted splicing probability positively correlates with splicing efficiency observed in the PTRE-seq library. Thresholds used to predict presence of moderate, strong, and very strong splicing are colored gray, purple, and pink, respectively. (B) Designs of evaluated 3’ UTR-focused MPRAs and predicted effect of cryptic splicing in these libraries. (C) Percentage of reporters predicted to be spliced from MPRAs described in (B). (D) Top enriched sequence motif in reporter transcripts with >0.60 splicing probability for each MPRA. Motif enrichment analysis was done using the XSTREME webserver^53^. Reporter transcripts with <0.30 splicing probability were used as matched controls. (E) Boxplots showing reporter expression as a function of predicted splicing probability. For clarity, data points outside the expression range (-6, 2) are not shown. (F) Opposing differences in RNA expression between paired ref and alt single-nucleotide variants in the *Griesemer* MPRA correlate with predicted splicing. alt > ref and ref > alt denote variants in which predicted splicing probability is 0.35 or greater between the ref and alt alleles. ref ≈ alt the difference in splicing probability is less than 0.35. Expression measurements from HEK293FT cells are shown^7^. For clarity, data points outside of the whiskers of the ref ≈ alt group are not shown. (G) Summary of RT-PCR validation of predicted spliced reporters from the *Griesemer* MPRA. See Figure S7 for raw data. (H) Example functional variant identified by the *Griesemer* MPRA that is explained by cryptic splicing. Bar plot shows mean relative expression measured by *Griesemer* with error bars denoting standard error^7^. SpliceAI probability is shown below along with the sequence of the predicted 3’ splice site (underlined). The variant is highlighted red. RT-PCR analysis of individually transfected reporter into HEK293T cells resolved by agarose gel is shown at right. For (E, F), box plot represents the distribution of splicing efficiencies for a group of reporters. Whiskers indicate the furthest datum that is 1.5*Q1 (upper) or 1.5*Q3 (lower). Statistical significance was tested using Mann-Whitney U rank test, where \**P* < 0.05, \*\**P* < 0.01, \*\*\**P* < 0.001.

We applied this spliceAI strategy to evaluate three other representative published MPRAs for which complete sequence information was readily available (Fig. 5B)^7–9^. Whereas PTRE-seq assayed synthetic 3’ UTRs, these other MPRAs assayed segments of natural human 3’ UTRs: the *Griesemer* library consists of 30,532 132-nt long segments containing putative functional 3’ UTR variants nominated by genome-wide association studies; the *Zhao* library consists of 2,828 highly conserved 200-nt long segments bearing diverse miRNA and RBP sites; and the *Siegel* library consists of 41,288 200-nt long segments bearing diverse AU-rich elements. Each of these MPRAs used GFP as the reporter gene, although placed in different plasmid architectures. These MPRAs used two different strategies to introduce reporters into cells: as exosomal plasmids (*Griesemer*) or integrated into chromatin using lentiviral vectors (*Zhao* and *Siegel*). Reporter expression for each MPRA was quantified by amplicon sequencing.

Our analysis indicates that these three MPRAs are also predicted to undergo cryptic splicing (Fig. 5C). In *Zhao* and *Siegal*, ∼10% of reporters have moderate to strong splicing probabilities. In *Griesemer*, ∼50% of reporters are predicted to undergo splicing, with 16% having very strong splicing probability (P>0.9). Similar to PTRE-seq, predicted splice donors were observed primarily at the same GFP site adjacent to the stop codon, but also occurred internally in the 3’ UTR (Fig. S6C). Splice acceptors were predicted to occur predominantly in 3’ UTR variable regions (Fig. S6C). The difference in predicted splicing susceptibility across MPRAs is explained in part by differences in reporter backbones that strengthen or weaken the cryptic GFP splice donor site (Fig S6D). Consistent with a critical role for U-rich sequences in driving splicing, motif analysis revealed strong enrichment of intronic U-rich motifs in predicted spliced reporters (Fig. 5D).

To test these predictions, we analyzed published expression measurements of the *Griesemer*, *Zhao*, and *Siegel* MPRAs for evidence of gene expression changes linked to splicing. Given the design of these MPRAs, we expected that splicing would disrupt the forward primer binding sites or excise regions used for reporter identification, resulting in sequencing dropout and apparent reduction in gene expression (Fig. 5B). Consistent with this expected impact, in the *Siegel* MPRA, reporters with predicted splicing probabilities above 0.6 and 0.9 exhibit 37% and 74% mean reduction in steady state expression, respectively, compared to low splicing probability reporters (p < 1.0× 10^−5^) (Fig. 5E). Similarly, in the *Zhao* MPRA, mean expression was reduced 43% and 90% for reporters with splicing probabilities above 0.6 and 0.9, respectively (p < 1.0× 10^−5^). The *Griesemer* MPRA exhibits significantly less dynamic range in reporter expression (Fig. 5E). Nevertheless, reporters with predicted splicing probabilities >0.9 exhibited 46% mean reduction in expression (p < 1.0× 10^−5^). This splicing-associated reduction in expression is greater than the repression induced by AREs, Pumilio binding sites, and miRNA binding sites (8% to 32% reduction) observed in the *Griesemer* study^7^. Together, these observations support that cryptic splicing is both present and has the potential to convolute interpretation of functional motifs identified by these MPRAs.

The study design of the *Griesemer* MPRA provides an additional opportunity to evaluate the impact of cryptic splicing on measured gene expression. This MPRA was designed to evaluate the functional effects of disease-linked 3’ UTR single-nucleotide polymorphosisms (SNPs) on gene expression^7^. Each assayed 3’ UTR segment was expressed in a pair of reporters bearing either the reference (ref) or alternative (alt) allele. Variants were then considered functional (termed a transcript abundance modulating variant, or tamVar) if they induced a significant difference in expression between the ref and alt reporters. 19 pairs of reporters in the *Griesemer* MPRA exhibit significant differences in predicted splicing probability (>0.35; Fig. 5F). For tamVars where the alt allele is predicted to be more highly spliced, the alt allele exhibits lower expression than the ref allele. Conversely, when the ref allele is predicted to be more highly spliced then expression of the ref allele is reduced (Fig. 5F). Thus, predicted splicing explains both increases and decreases in expression caused by SNPs. An additional 266 tamVars were predicted to have high splicing probabilities (>0.9) in both alleles, indicating that cryptic splicing likely convolutes these tamVar measurements. Conservatively, our analysis suggests that 285 out of 2,368 SNPs (12%) identified as functional by *Greisemer* et al may be impacted by cryptic splicing.

To independently validate our spliceAI predictions, we synthesized 20 reporters from the *Griesemer* MPRA comprising 10 ref/alt allele pairs identified as functional tamVars. These tamVars represent SNPs associated with prostate cancer, schizophrenia, anxiety disorder, and human evolution^7^. 2 pairs of reporters were predicted to be both unspliced, 5 were predicted to be both highly spliced, and 3 were predicted to be differentially spliced (one high, one low) (Fig. 5G, S7). Reporters were individually transfected into HEK293 cells and splicing was assessed using semi-quantitative RT-PCR and sequencing. 12 out of 13 predicted highly spliced reporters demonstrated splicing, corresponding to a positive predictive value of 92%, whereas all 7 low splicing probability reporters were unspliced (Fig. 5G, S7A-C). For example, SpliceAI predicted that the rs13004845 tamVar contains a strong 3’ AG splice site in the ref allele which is abolished by a G-to-A substitution in the alt allele (Fig. 5H). Our experiments confirmed that the ref allele was spliced with 57% efficiency, whereas the alt allele is unspliced, consistent with the 28% decrease in ref allele expression measured by the *Griesemer* MPRA. Similarly, variants rs5756095 and rs140761234 create and weaken predicted splicing acceptor sites, respectively, inducing differential splicing that explains the changes in expression measured by MPRA (Fig. S7B). 3 of the reporter pairs predicted to be highly spliced in both ref and alt alleles also exhibited significant differences in splicing efficiencies consistent with the differential expressions observed by MPRA (Fig. S7C). Overall, these results emphasize the ability of cryptic splicing to result in spurious conclusions across diverse MPRA studies, impacting interpretation of gene regulatory mechanisms and misclassification of human SNPs.

## DISCUSSION

Cryptic splicing is a well-known phenomenon^4^, but one that it is generally assumed to be rare. In this study, we report that cryptic splicing likely affects between 10% and 50% of GFP reporter genes bearing functional 3’ UTRs. Using resequencing and reanalysis of the PTRE-seq MPRA, we establish that cryptic splicing is predominantly determined by the presence and arrangement of intronic AU-rich elements that both potentiate splicing and dictate 5’ and 3’ splice site choice. We further show that this cryptic splicing has diverse impacts on reporter assay measurements, with potential to both upregulate and downregulate apparent gene expression depending on splice site locations and measurement strategy. These findings have important implications for the future design and interpretation of MPRAs and reveal new insights into the sequence codes governing splicing activation and splice site choice^23^.

A major driver of splicing in the PTRE-seq library is an undocumented cryptic splice donor in the GFP coding sequence. While relatively weak compared to endogenous donors, this site can still drive efficient splicing into the 3’ UTR given the presence of U-rich sequences and viable downstream splice acceptors (Fig. 2). Our analysis of other MPRAs supports that this splicing site is broadly used in other reporter contexts, including MPRAs that were expressed from lentiviral integrated transgenes. Since this donor site overlaps the stop codon of GFP, the resulting GFP protein product contained a C-terminal extension and exhibited lowered stability (Fig. S2)^28^, emphasizing that cryptic splicing can impact gene expression at multiple levels. We show that this GFP-driven splicing can be suppressed using synonymous recoding of the -4 and -2 C-terminal codons (Fig. 2). We suggest that these synonymous mutations should be broadly incorporated into fluorescent reporter genes to mitigate undesirable splicing. However, we emphasize that efficient splicing also occurs from donors within 3’ UTRs, indicating that there is unlikely to be a universal solution for eliminating cryptic splicing.

Our results revealed an unexpectedly central role of U-rich elements as potent activators of cryptic splicing. In PTRE-seq, AREs were capable of activating highly efficient splicing from otherwise weak 5’ and 3’ splice sites. This ability to activate splicing includes but is not limited to the ability of AREs to serve as a polypyrimidine tract; for example, AREs are able to activate splicing 30 nucleotides downstream at the weak “spacer” acceptor site (Fig. 3C). A limitation of our study is that PTRE-seq only assays a single ARE (U-rich) sequence, albeit in many different contexts. Defining a more generalized understanding of the U-rich sequence features responsible for splicing activation is an important topic for future studies. Nevertheless, we show that U-rich motifs are associated with cryptic splicing signatures in other MPRAs (Fig. 5D). Our results are also consistent with the well-documented ability of U-rich tracts in *Alu* and other transposable elements to activate weak splice sites and drive evolution of new exons^22,38^. In splicing of evolved (non-cryptic) exons, intronic U-rich elements are strongly associated with high splicing efficiency and with activation of proximal alternative 5’ splice sites^39,40^. Collectively, these data support a model in which U-rich elements can function as principal splicing stimuli.

We also found that AREs exert a powerful position-dependent influence on both 5’ donor and 3’ acceptor splice site choice (Fig. 3). Most notably, placement of AREs in exonic locations relative to potential donor and acceptor sites (5’ and 3’ of donor sites, respectively) is sufficient to completely suppress splice sites that are efficiently spliced in other contexts. A number of alternative splicing factors are known to bind U-rich sequences and exhibit position-dependent effects on splice site selection^22,39,41,42^. However, a dominant role for exonic U-rich sequences in prohibiting splice site selection has not been previously appreciated. Further investigation is needed to define the mechanisms behind the positional effects of AREs. Overall, these findings suggest that U-rich sequences may play broader roles in defining intronic architecture than previously appreciated.

Our observation that cryptic splicing is activated by only relatively simple U-rich motifs also has implications for understanding ubiquitous mis-splicing events in disease. U-rich motifs with appropriately spaced downstream AG sites that could serve as cryptic splice acceptors are prevalent throughout the transcriptome. For example, we find that many natural 3’ UTR sequence elements serve as efficient splice acceptors in MPRA contexts (Fig. 5). These natural 3’ UTR sequences go unspliced in their endogenous contexts, implying that mechanisms normally suppress splicing at these 3’ UTR sites and that dysregulation of these mechanisms could be sufficient to activate cryptic splicing^22,43^. Indeed, a recent study reported widespread upregulation of cryptic splicing in 3’ UTRs in cancer, with these splicing events also exhibiting a strong association with U-rich elements^44^. Some of the disease-associated mutations that activate cryptic splicing in MPRA contexts (Fig. 5) may also be “functional” in endogenous contexts, inducing cryptic splicing that impacts gene expression rather than the originally assumed post-transcriptional mechanism.

In the context of MPRAs, cryptic splicing has complex impacts on reporter expression, including reducing apparent expression due to sequencing dropout, altering expression via removal of regulatory elements, and enhancing the production of spliced reporters (Fig. 4B, 5B). The 2-4 fold enhanced production we observed in PTRE-seq is consistent with prior measurements of intron-mediated enhancement of transcription^30–33^. However, alternative mechanisms such as facilitating mRNA export or promoting polyadenylation are also possible^45,46^. Interestingly, our analysis suggests that enhancement of production only emerges when reporters are spliced with >90% efficiency (Fig. 4C). Studies of the likely related phenomenon of exon-mediated activation of transcription starts (EMATS) also observed strong threshold effects, with only introns spliced in at >95% frequency inducing strong EMATS^33^. The mechanisms behind splicing-induced production enhancement and the threshold effects observed in our analyses require further validation, but are reminiscent of threshold effects observed during phase-separation processes and we speculate may be linked to partitioning into transcriptional condensates^47,48^.

Our re-analysis of published MPRAs revealed that cryptic splicing and other artifacts (RT recombination, barcode dropout) can significantly impact MPRA conclusions. In the PTRE-seq MPRA, ARE-induced splicing led to the erroneous conclusion that AREs can stabilize mRNAs and promote translation depending on their 3’ UTR context. Once splicing is accounted for, we find that AREs uniformly reduce mRNA expression, consistent with the prevailing model of ARE function^8,34,35^. In other 3’ UTR-focused MPRAs, we provide evidence that splicing-induced sequencing dropout results in anomalously low mRNA expression, comparable to or greater than the reductions in expression induced by bona fide post-transcriptional regulatory motifs. Because MPRAs are often analyzed with the goal of identifying sequence elements that convey strong regulatory effects, these cryptically spliced sequences are particularly liable to be identified as functionally interesting (Fig. 5). Indeed, we show that cryptic splicing likely impacts more than 10% of functional SNPs identified by a recent MPRA designed to assess human disease-associated variants^7^. Nevertheless, it is important to emphasize that the majority of MPRA measurements remain valid, though they should be interpreted with appropriate caution.

In conclusion, cryptic splicing is unexpectedly common in MPRAs due both to MPRA design features and the apparent ease at which splicing is activated in transgenes. As a precautionary measure, we recommend utilizing a splicing prediction algorithm such as SpliceAI to identify potential cryptic splice sites during MPRA design^36^. In addition, diverse sequencing-related artifacts such as reverse transcription errors reduce usability of MPRA data. Therefore, we recommend using sequencing strategies that can reliably identify splicing artifacts and other experimental errors, such as performing full-length sequencing. These quality control steps will ensure more robust and reliable MPRA experimental outcomes and interpretation.

## METHODS

### PTRE-seq library design

Construction of the pooled PTRE-seq plasmid library was described previously^24^. In brief, a pooled library of synthetic 3’ UTRs was cloned downstream of eGFP in the pCDNA5/FRT/TO plasmid (Addgene 19444). Each 3’ UTR consisted of a 19-nucleotide fixed spacer, a 132-nucleotide variable regulatory array, a 20-nucleotide spacer, a 9-nucleotide identifying barcode, and a 225-nucleotide fixed sequence terminating with the bGH polyA sequence. Each of the variable regulatory arrays consists of 4 33-nucleotide long regulatory modules encoding a combination of (a) “blank” control sequences, (b) let-7 miRNA binding sites, (c) Pumilio protein recognition sites, (d) Smaug protein recognition sites, and (e) AU-rich elements (Fig. 1). Each regulatory array is present in 10 copies, each bearing a different barcode, which function as internal replicates. In addition, the library contains 40 additional copies of the control sequence (4 “blanks”), 50 copies of a low expression control (4×let-7 perfect complement), and a series of constructs containing natural and synthetic let-7 sites.

### Cell culture and transfection of PTRE-seq libraries

HeLa (Baylor College of Medicine Tissue Culture Core), HEK293 (R78007, ATCC), SH-S5Y5 (CRL-2266, ATCC), U87 MG (HTB-14, ATCC), and T-REx TM-293 cells (R71007, Thermo Fisher) cells were grown in DMEM (Gibco) supplemented with 10% FBS (Gibco), 1x Penicillin streptomycin and glutamine (Gibco) and 1x MEM non-essential amino acids (Gibco). Plasmid libraries were transfected using the Neon Transfection System (Invitrogen) per manufacturer protocol. For each transfection, 2.5 x 10^6^ cells were electroporated with 8 μg of the PTRE-seq library. RNA was isolated and purified 40 hours post-transfection via RNeasy mini kit (Qiagen), treated with Turbo DNase (Invitrogen, AM2238) and purified using Mag-Bind TotalPure NGS beads (Omega Bio-tek). Cells were regularly tested and confirmed to be mycoplasma free.

Transfection of individual PTRE-seq plasmids and site-directed mutants were performed using Effectene (Promega) or X-tremeGENE™ 9 DNA Transfection Reagent (Roche), respectively. Cells were transfected in a 6-well plate with 1000 μg plasmid, then split 24 hr later into two separate 6-well plates. After 40 hours, RNA was isolated using RNeasy kit (Qiagen), treated with Turbo DNase (Invitrogen) and purified using Mag-Bind TotalPure NGS beads (Omega Bio-tek).

### Semi-quantitative PCR of individual PTRE-seq reporters

2 μg total RNA from individual transfections was reverse transcribed using Superscript II reverse transcriptase (Invitrogen) using RE_Amp_RT primer or Junction_R (Table S1). cDNA product was PCR amplified (Q5, NEB; 98 °C for 30 s, 35 cycles: 98 °C for 10 s, 71 °C for 20 s, 72 °C for 30 s, and 72 °C for 2 min) using GFP_Amp_F_seq and RE_Amp_R_seq or Junction_R primers (Table S1) and visualized on 1.5% agarose gel stained with ethidium bromide.

### RNA and DNA sequencing of PTRE-seq libraries

RNA sequencing libraries from pooled library transfections were prepared from 2 μg total RNA. Reverse transcription was performed using Superscript II reverse transcriptase (Invitrogen) or MarathonRT (Kerafast) using the RE_Amp_RT primer. We adapted a previously described 2-step PCR protocol ^49^ to amplify the 3’ UTR and attach Illumina adaptors with indexes. To minimize PCR chimeric artifacts, three emulsion PCR (Micellula DNA Emulsion & Purification Kit, EURx) reactions were prepared independently and then pooled for each experiment. For step 1 PCR, 1 μL of the purified cDNA product was amplified using GFP_Amp_F_seq and RE_Amp_R_seq primers (98 °C for 30 s, 20 cycles: 98 °C for 10 s, 71 °C for 20 s, 72 °C for 30 s, and 72 °C for 2 min). For step 2 PCR, 100 pg of PCR step 1 product was amplified using step 2 universal adaptor primers (98 °C for 30 s, 15 cycles: 98 °C for 10 s, 71 °C for 20 s, 72 °C for 30 s, and 72 °C for 2 min). Matching DNA sequencing libraries from the DNA plasmid were prepared by inputting 1 μL of plasmid into the same two-step PCR protocol. Libraries were sequenced using either an Illumina MiSeq (v2 chemistry, 2 x 250 bp; or v3 chemistry, 2 x 300 bp) or an Illumina NovaSeq 6000 (SP flowcell 2 x 250 bp).

### Western blot analysis

Protein lysates were collected from cells transfected with individual reporters (cell lysis solution, Cell Signaling Technology), resolved on a 4–12% Bis-Tris gel (Bio-Rad), and transferred onto Immuno-Blot PVDF (Bio-Rad). After blocking (5% milk in 1x PBS with 1% Tween, 1 hour minimum), proteins were incubated with primary antibodies against GFP (Clonetech, 632381) at 1:5000 dilution and β-actin-HRP (Cell Signaling, 12261) at 1:1000 dilution for 1 hour at room temperature or overnight at 4°C. Following washing with 1x PBST, proteins were incubated with anti-mouse secondary antibody coupled to horse-radish peroxidase (Cell Signaling, 12262) at 1:10000 dilution. After washing with 1x PBST, imaging was conducted using a Bio-Rad ChemiDoc XRS System.

### Mutation of GFP splice donor sites in PTRE-seq reporters

Site-directed mutagenesis was used to introduce synonymous mutations that ablate the GFP splice donor sites in the ASAS and AALB reporters. PCR was performed following manufacturers instructions (Q5; NEB) using M1_For and M1_Rev primers for mutant 1 and M2_For and M2_Rev primers for mutant 2 (Table S1). Plasmids were transformed into *E. coli*, purified with Qiagen Plasmid Midi Kit (Qiagen), and confirmed via sanger sequencing.

### Barcode mapping and splice isoform quantification

Sequencing data were processed by first merging paired-end reads using BBMerge with default parameters^50^. The pooled MPRA library was then demultiplexed based on internal barcodes using a custom Python script. The edit distance was calculated between each sequencing read and all 6500 barcodes, which include 4 flanking nucleotides upstream and downstream (align sequence = <u>CGAG</u> + 9-nt barcode + <u>GGTA</u>). Reads were assigned to a reporter if the edit distance was 1 or less. BBMap^51^ was then used to align assigned reads to the reference sequence of the reporter using default parameters.

Following demultiplexing and mapping, each output bam file was analyzed using a custom Python script to classify reads into one of the five categories:

i. RT recombination: we observed a number reads featuring a reverse complement of one of the spacer regions accompanied by neighboring indels (Supplementary Fig. S1E). We ascribed these reads to premature template switching and recombination during reverse transcription. Reads were assigned as “RT recombination” if the spacer sequence was immediately preceded by the “CGG” trinucleotide instead of the expected GCC.
ii. Chimeric: PCR chimeras can arise from annealing of incomplete extension products, which can result in the barcode from one reporter being appended to a different reporter. After barcode demultiplexing, chimeric reads will poorly align to their expected reference sequence, resulting in numerous indels. Reads were assigned “chimeric” if they featured more than two indels in the regulatory array region.
iii. Spliced: reads were assigned as spliced if they (a) contained a deletion flanked by two matching sequences (b) featured GT and AG dinucleotides at the 5’ and 3’ end of the deletion, respectively, and (c) featured overall lengths within 5 nts of the expected length for the spliced isoform.
iv. Full length: reads were assigned as full length if they (a) possessed no more than 3 mismatches when aligned against the reference sequence and (b) featured lengths within 5 nts of the expected 456 nt full amplicon length.
v. Ambiguous: reads were assigned as ambiguous if they featured more than 3 mismatches relative to the expected reference sequence, or featured a long indel without an GT-AG junction.

The splicing efficiency for each barcode was calculated as:

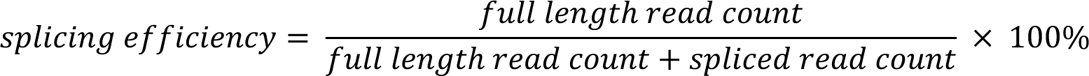

Each regulatory array is present in 10 copies in the PTRE-seq library, each with a different barcode, resulting in 10 possible splicing efficiency measurements per biological replicate. In order to compute a barcode-specific splicing efficiency, we required a minimum of 10 total spliced and full-length counts. Final splicing efficiency measurements were computed as the median of all measurements passing this minimum count filter (a maximum of 20 measurements: 10 internal replicates x 2 biological replicates).

### Linear regression model

We adapted a linear regression strategy developed in the original PTRE-seq study to model reporter expression as a function of 3’ UTR regulatory elements^24^. The variables of this model include the identity of the regulatory element at each of the four positions and their pairwise interaction. The parameters of the model represent the impact of each variable on reporter expression relative to the “blank” control. We used the Generalized Linear Model (*glm*) from the Statsmodels python package with the following formula:

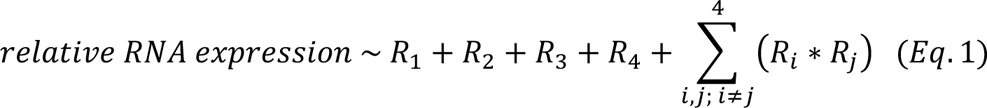

Where *Ri* denotes the regulatory element at each 3’ UTR regulatory position. To avoid the confounding effect of splicing, we only fit the model to the 476 reporters exhibiting splicing efficiencies below 1%. The resulting parameters were used to estimate expression of the full-length isoform of the remaining 149 reporters. Low expression controls and reporters with natural and synthetic let-7 sites were excluded from analysis. The model was fit using expression measurements from the original PTRE-seq study^24^.

### Gene expression modelling and transcription rate estimation

At steady state, mRNA expression (*E_tot_*) of each spliced reporter can be expressed as

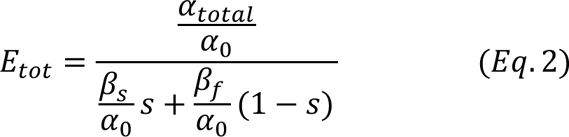

where *α_total_* is the combined production rate of the spliced and full-length reporter mRNAs, *α*_0_ is the production rate of unspliced reporters, *β_s_* and *β_f_* is the degradation rate for the spliced isoform and full-length isoform respectively, and *s* is the splicing efficiency.

The stability of the full-length isoform, 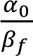, is estimated from the linear regression model described in Eq. 1. To obtain the stability of the spliced isoform, 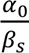, we rearrange Eq. 2 to the form:

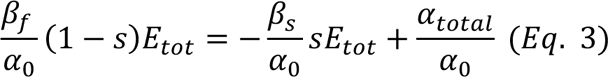

Reporters that are spliced using the same 5’ and 3’ splice sites will have identical spliced isoforms and hence identical stabilities. Thus, for families of reporters with identical spliced isoforms, 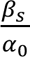 can be obtained as the slope from the regression of 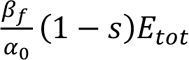 against *sE_tot_*. We identified two families of reporters that have sufficient variation in *sE_tot_* to permit this regression: GFP_B4, where reporters feature splicing from the primary GFP donor to a 4^th^ position B module, and GFP_P4, where reporters are spliced at the GFP donor to a 4^th^ position PRE acceptor. For each family, we tested fitting the model using reporters within different ranges of splicing efficiencies (10-70%, 10-80%, and 10-90%) to assess the robustness of the model. Fitting was done with the *glm* function from the Statsmodels Python package using expression measurements from the original PTRE-seq study^24^ and splicing efficiency measurements made in the current study. Fitting the model using expression measurements from the current study yielded similar results. The RE-excision model is then obtained from Eq. 2 under the assumption of no production changes (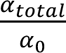 = 1). To estimate production rate, we rearrange Eq. 2 to solve for 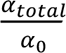

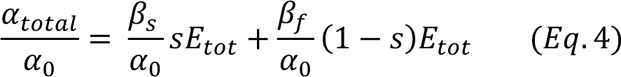

### Prediction of cryptic splicing in published MPRAs

We adapted SpliceAI to predict splicing probabilities for the reporter sequences following the SpliceAI documentation^36^. For the PTRE-seq library, the input sequence for each reporter is a 1172-nt transcript, which begins with the eGFP coding sequence and ends at the poly-A signal. The sequence was padded with “N” until the length was 10,000 nucleotides. SpliceAI outputs the probability that each position on the transcript is a donor or acceptor. The product of the donor and downstream acceptor with the highest probability is calculated as splice probability and is used to predict whether a reporter is spliced or not. We explored other potential total splicing scores, including considering only donor or acceptor probabilities, sum of both probabilities, and Maximum Entropy scores^52^; these scores performed comparably or worse than the product at predicting PTRE-seq splicing efficiencies.

For other MPRAs (*Griesemer*, *Zhao*, and *Siegel*), sequence information of the inserted regions and the backbone plasmids was retrieved directly from electronic supplementary data or from the authors. The transcript sequence for each reporter was then reconstructed beginning with the eGFP coding sequence and ending at the poly-A signal. Barcode sequences were added as 6 N(s) for *Griesemer* MPRA and 8 N(s) for *Siegel* and *Zhao* MPRA. Splicing probabilities were then computed identically as for PTRE-seq.

### Motif enrichment analysis

XSTREME was used to identify enriched motifs in predicted spliced reporters^53^ from the *Griesemer*, *Siegel*, and *Zhao* MPRAs. The sequences of reporters above 0.60 splicing probability were uploaded in fasta format to the XSTREME webserver (https://meme-suite.org/meme/tools/xstreme). Sequences of reporters below 0.30 splicing probability were uploaded as controls. All parameters were set to defaults. Motifs shown in Figure 5D correspond to the most significantly enriched motif for each MPRA. The large majority of other significantly enriched sequences were also U-rich motifs.

### Validation of splicing in *Griesemer* MPRA reporters

10 pairs of alt/ref reporters (20 overall) were selected from the *Griesemer* MPRA for validation^7^: as negative controls, 2 pairs where both alleles were predicted to be not spliced; 3 pairs where the alleles were predicted to be differentially spliced; and 5 pairs where both alleles were predicted to be spliced with high probabilities. Synthesis and cloning were done by the Genetic Design and Engineering Center at Rice University. DNA sequences containing 3’ UTR segments, random 8 nt barcodes, and flanking adaptors matching the final assembled inserts in *Griesemer* were ordered as eBlocks (IDT) (Table S2). Each eBlock was amplified by PCR using the Griesemer_Amp_F and Griesemer_Amp_R primers (Table S1) and then cloned by Gibson Assembly (NEB) into BmtI/XbaI (NEB) digested pmirGLO:Δluc::gfp ΔAmp^R^::Kan^R^ vector (gift from Pardis Sabeti; Addgene #176640). The assembled vectors were transformed into 10-beta *E. coli* (NEB) by electroporation, expanded in LB broth supplemented with 50 μg/mL of kanamycin, and plasmids purified via Qiaprep Spin Miniprep Kit (Qiagen). Plasmid sequences were verified by Sanger sequencing.

Reporters were transfected using Lipofectamine 2000 (Thermofisher) according to the protocol published by Griesemer *et al* ^7^. HEK293T cells were grown in a 6-well plate until 70% confluency. 3 *μ*g of DNA was combined with 9 *μ*L of Lipofectamine 2000 in a 300 *μ*L DMEM solution and added to a well. After 24 hr, RNA was harvested using RNeasy kit. cDNA was prepared using Griesemer_R primer and SuperScript II reverse transcriptase per manufacturer protocol. 4 μL of the purified cDNA product was PCR amplified (98°C for 30s, 35 cycles: 98°C for 10s, 68°C for 20s, 72°C for 30s, and 72°C for 2 min) using Griesemer_F and Griesemer_R primers (Table S1). PCR product was visualized on 1.5% agarose gel. Splicing was also validated using sequencing. Sequencing libraries were prepared by inputting 2 μL of cDNA into step 1 PCR (98 °C for 30 s, 15 cycles: 98 °C for 10 s, 68 °C for 20 s, 72 °C for 30 s, and 72 °C for 2 min) using GFP_Amp_F_seq and RE_Amp_R_seq primers (Table S1). For step 2 PCR, 10 pg of PCR step 1 product was amplified using step 2 universal adaptor primers (98 °C for 30 s, 10 cycles: 98 °C for 10 s, 68 °C for 20 s, 72 °C for 30 s, and 72 °C for 2 min). Sequencing was done using an Illumina MiSeq (v2 chemistry; 2 x 250).

### Data availability

Analysis scripts and Jupyter Notebooks are available at the Github repository under MIT license (https://github.com/MustoeLab/Publications/Dao_MPRAsplicing/). Sequencing data is available at the Sequence Read Archive under accession code SRP509815, and project details can be found in the NCBI BioProject under accession code PRJNA1116243.

## Acknowledgements

We thank D. Griesemer, J. Xue, P. Sabeti, W. Zhao, D. Siegel, and D. Erle for sharing sequences and data from their studies, and K. Cottrell for helpful discussions. We thank T. Westbrook for gifting us the HEK293T cell line, the Genetic Design and Engineering Center at Rice University for cloning services, and C. Chen and M. Goodell for the use of their electroporator.

This work was supported by a CPRIT scholar award to A.M.M. by the Cancer Prevention and Research Institute of Texas (RR190054). S.D. and C.F.J. are supported by NIH GM112824 and Chan Zuckerberg Neurodegeneration Initiative Award. The Genomic and RNA Profiling Core at Baylor College of Medicine is supported by P30 Cancer Center Support Grant (NCI-CA125123). The Genetic Design and Engineering at Rice university is supported by CPRIT RP210116.

