## Supporting Information for "U-rich elements drive pervasive cryptic splicing in 3’ UTR massively parallel reporter assays"

**Figure S1**

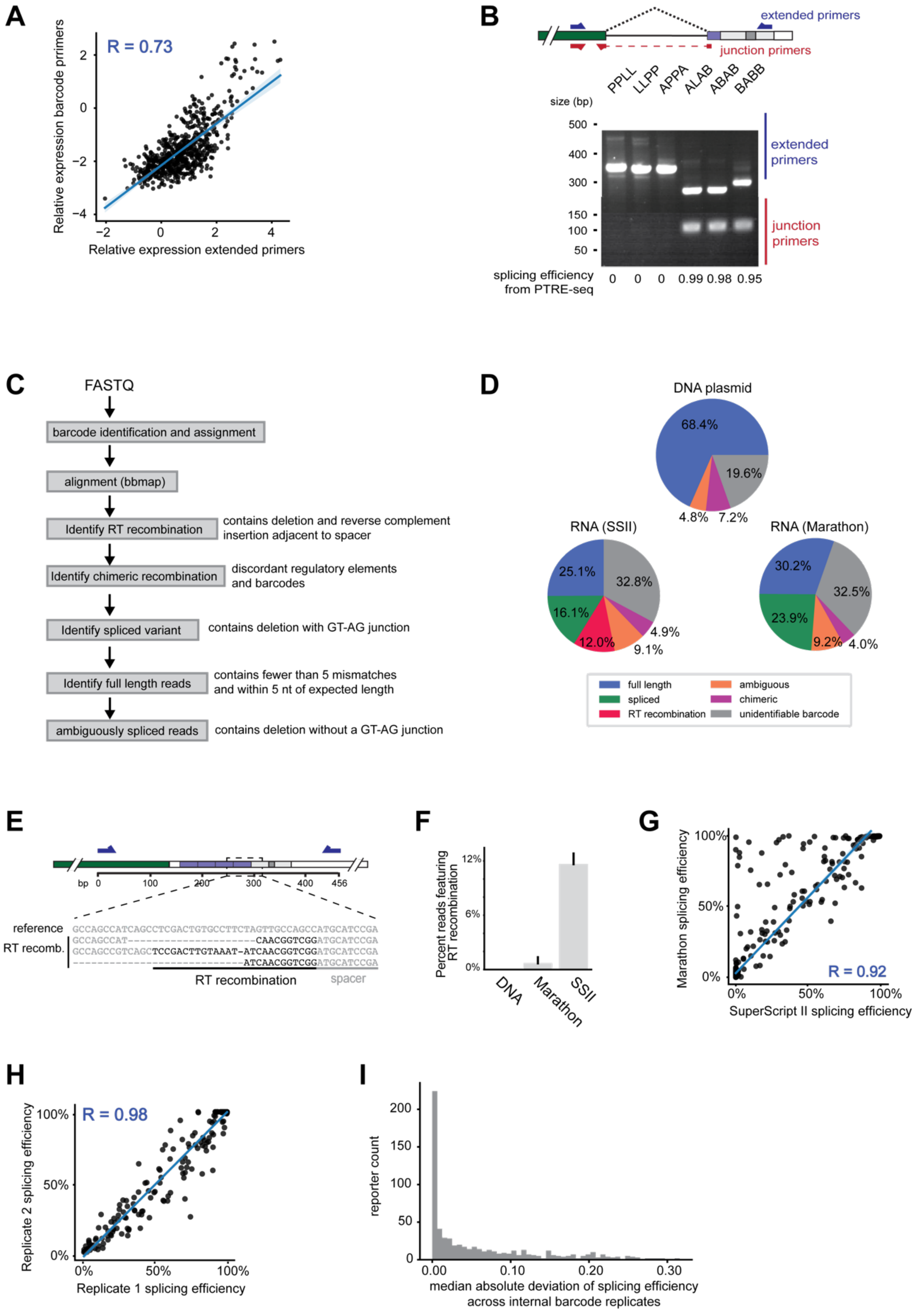
A
R = 0.73
Relative expression barcode primers
Relative expression extended primers
B
extended primers
junction primers
PPLL
LLPP
APPA
ALAB
ABAB
BABB
size (bp)
500
400
300
150
100
50
extended primers
junction primers
splicing efficiency from PTRE-seq
0 0 0 0.99 0.98 0.95
C
FASTQ
barcode identification and assignment
alignment (bbmap)
Identify RT recombination
contains deletion and reverse complement insertion adjacent to spacer
Identify chimeric recombination
discordant regulatory elements and barcodes
Identify spliced variant
contains deletion with GT-AG junction
Identify full length reads
contains fewer than 5 mismatches and within 5 nt of expected length
ambiguously spliced reads
contains deletion without a GT-AG junction
D
DNA plasmid
68.4%
19.6%
4.8%
7.2%
RNA (SSII)
25.1%
32.8%
16.1%
12.0%
4.9%
9.1%
RNA (Marathon)
30.2%
32.5%
23.9%
9.2%
4.0%
full length
spliced
RT recombination
ambiguous
chimeric
unidentifiable barcode
E
bp 0 100 200 300 400 456
reference GCCAGCCATCAGCCTCGACTGTGCCTTCTAGTTGCCAGCCATGCATCCGA
RT recomb. GCCAGCCAT-------------------------CAACGGTCGGATGCATCCGA
GCCAGCCGTCAGCTCCGACTTGTAAAT-ATCAACGGTCGGATGCATCCGA
-----------------------------ATCAACGGTCGGATGCATCCGA
RT recombination
spacer
F
Percent reads featuring RT recombination
12%
6%
0%
DNA
Marathon
SSII
G
Marathon splicing efficiency
SuperScript II splicing efficiency
R = 0.92
H
Replicate 2 splicing efficiency
Replicate 1 splicing efficiency
R = 0.98
I
reporter count
median absolute deviation of splicing efficiency across internal barcode replicates

**Figure S1. Quality control and quantification of splicing and library preparation artifacts in the PTRE-seq library.** (A) Comparison of RNA expression measurements obtained from extended primers (this study) compared to barcode primers (ref. 24). (B) RT-PCR validation of splicing on individually transfected reporters. Reporters were transfected into HeLa cells and RT-PCR products were resolved on agarose gels using extended and junction primers. Splicing efficiency measured from sequencing of the PTRE-seq MPRA is shown at bottom. (C) Analysis pipeline used for detection and quantification of splicing and other artifacts in the PTRE-seq library. (D) Quantification of read categories from sequencing of the PTRE-seq libraries prepared from plasmid DNA, or RNA reverse-transcribed with SuperScript II or MarathonRT. Label shows the mean percentage for each category over two biological replicates. (E) Example reads showing putative RT recombination artifacts. (F) Percent of “RT recombination” artifacts detected in DNA plasmids and in RNA sequencing library prepared from Superscript II or Marathon. Error bars show standard deviation over two biological replicates. (G) Comparison of splicing efficiency measurements obtained from libraries prepared using Superscript II versus MarathonRT. (H) Comparison of splicing efficiency measurements made across biological replicates in HeLa cells. (I) Median absolute deviation of splicing efficiency computed across alternative barcodes for each 3’ UTR. In (A, G, H), line of best fit and Pearson correlation coefficient is shown in blue.

### Figure S2

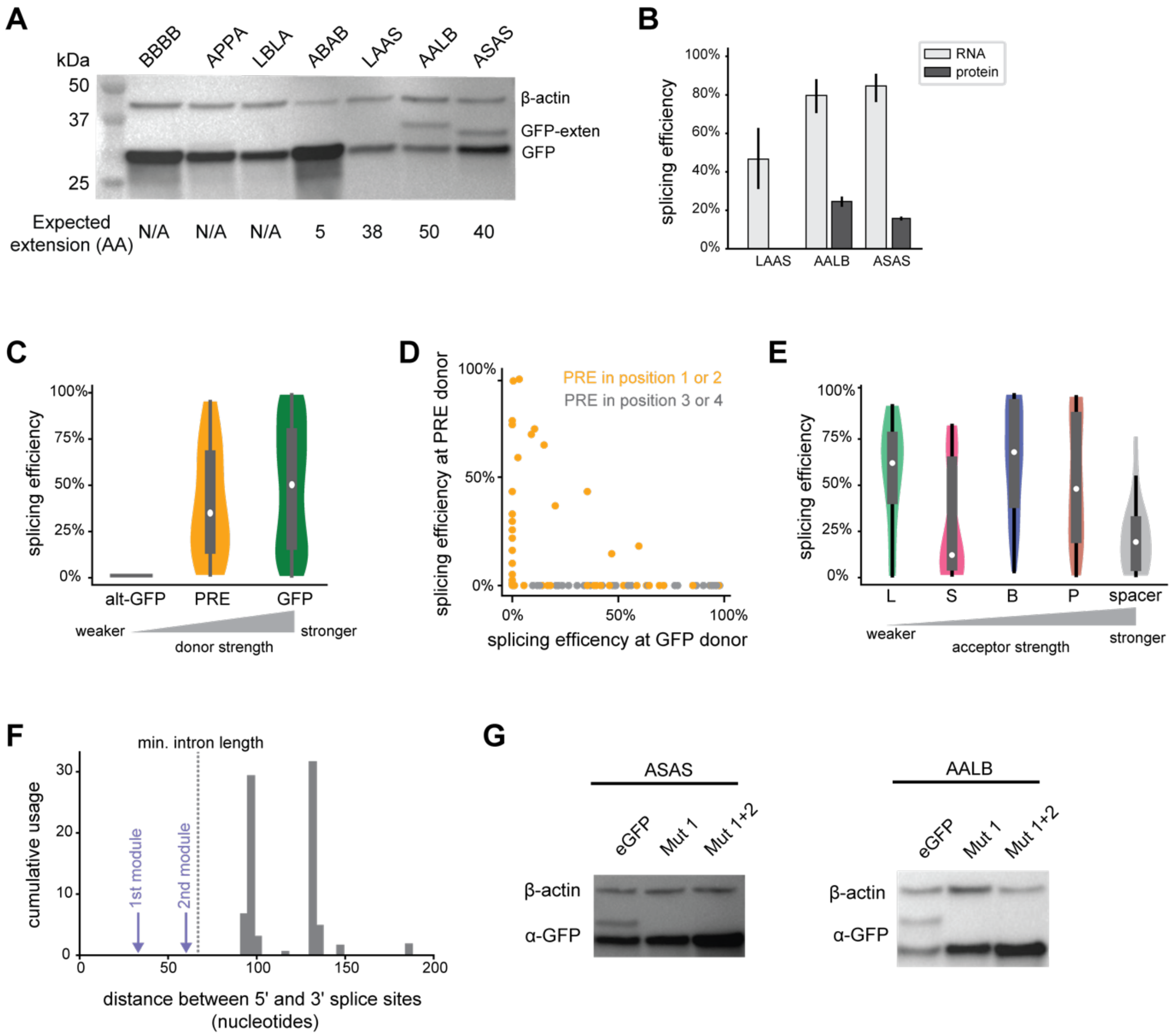


**Figure S2. Validation and analysis of cryptic splicing donor and acceptor sites in PTRE-seq library.** (A) Western blot analysis of GFP products from transfections of individual reporters. β-actin is shown as loading control. Predicted amino acid (AA) extension based on distance to the nearest in-frame stop codon is shown at bottom. (B) Comparison of spliced fraction measured by MPRA sequencing and western blot from spliced reporters in (A). The ABAB reporter was excluded from this analysis because its spliced versus full-length isoform cannot be distinguished via western blot. Error bars show standard deviation from 3 biological replicates. (C) Distribution of splicing efficiencies exhibited by different PTRE-seq splice donors. (D) Scatterplot of PRE donor versus GFP donor usage in PRE-containing reporters. (E) Distribution of splicing efficiencies observed at different splice acceptors in the PTRE-seq library. (F) Distributions of intron lengths observed in the PTRE-seq library, weighted by cumulative usage. Dashed line indicates minimum intron length identified from previous studies[29]. Purple arrows indicate intron lengths if 1$^{st}$ or 2$^{nd}$ module sites were used as splice acceptors. (G) Western blot analysis of ASAS and AALB reporters where GFP was synonymously recoded to abolish the 5' cryptic splice site. Matching RT-PCR data are shown in Figure 2F. For (C) and (E), violins illustrate the probability density, boxplots denote quartiles, white dots represent the medians, and whiskers represent the min and max for each distribution.

### Figure S3

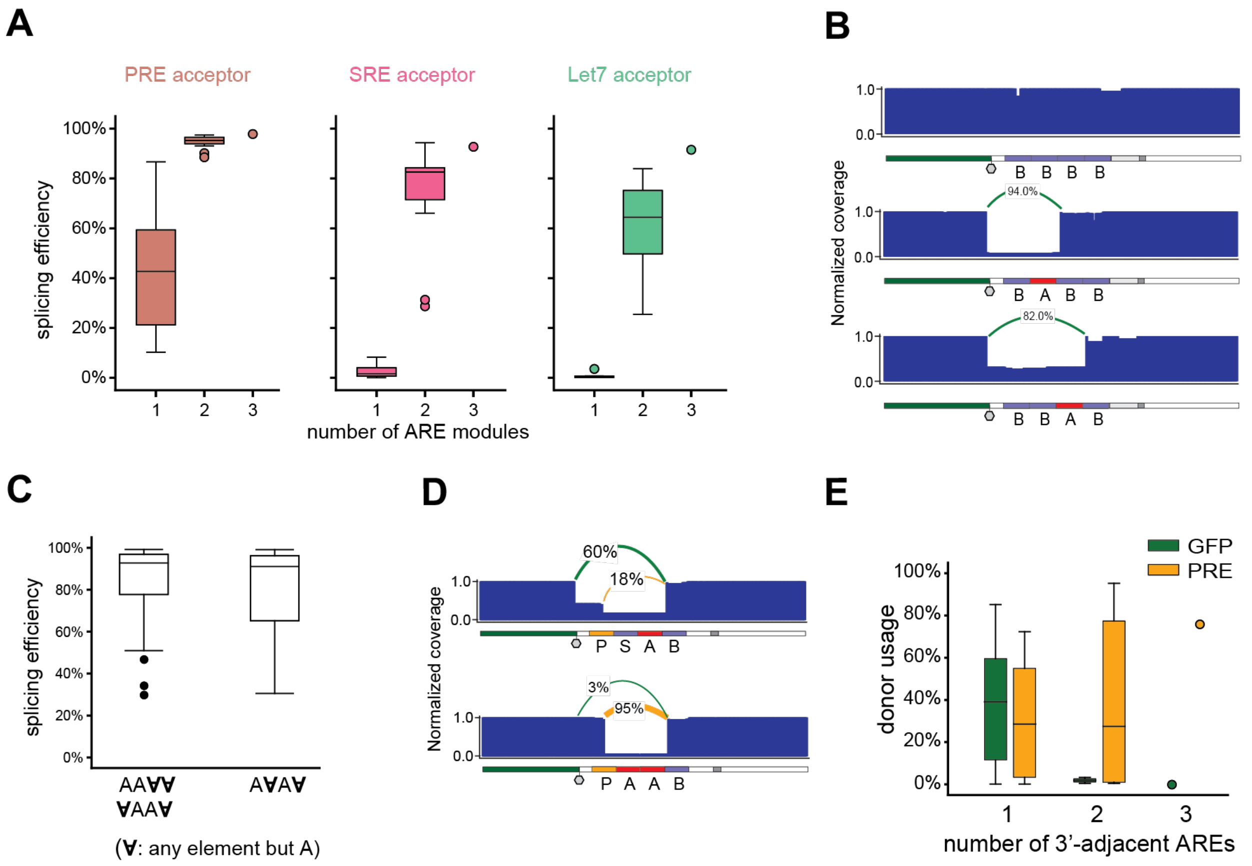


**Figure S3. AREs are position-dependent regulators of cryptic splicing.** (A) Relationship between the number of intronic AREs and splicing efficiency for different splice acceptors. (B) Example read coverage tracts showing a single ARE module is sufficient to activate highly efficient splicing at the B acceptor. (C) Comparison of splicing efficiency between reporters containing two AREs that are contiguous versus non-contiguous. (D) Example read coverage tracts illustrating ability of adjacent AREs to bias selection of PRE versus GFP donors. (E) Comparison of splice donor usage in reporters with increasing numbers of ARE modules downstream of the PRE donor. For (A, C, E), each box plot represents the distribution of spliced fraction for a group of reporters. Whiskers indicate the furthest datum that is 1.5*Q1 (upper) or 1.5*Q3 (lower).

### Figure S4

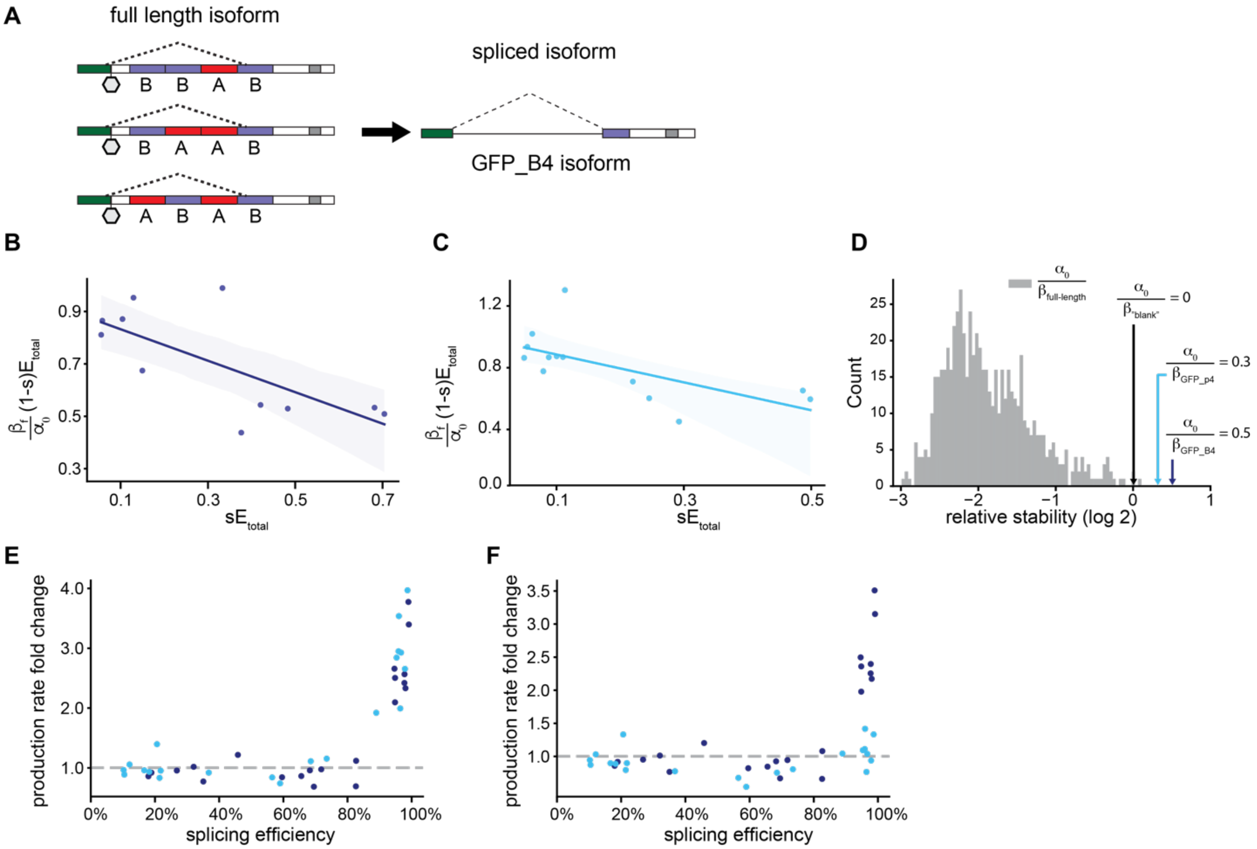


**Figure S4. Modelling the impact of splicing on mRNA expression using RE-excision and production enhancement models.** (A) Schematic showing a family of reporters with different pre-mRNA sequences that are spliced at the same donor and acceptor sites, resulting in the same GFP_B4 spliced isoform. (B) Fit of Equation 3 to the GFP_B4 family of reporters using reporters spliced from 10% to 80% efficiencies. The stability of the GFP_B4 isoform ($\frac{\beta_s}{\alpha_0}$) is obtained from the slope of the linear regression model. Translucent band shows the 95% confidence interval for the regression line. (C) same as (B), but for the GFP_P4 family of reporters. (D) Estimated stability coefficients for the spliced isoforms GFP_B4 and GFP_P4 compared to stability coefficients of full-length reporters. $\frac{\beta_s}{\alpha_0}$ for GFP_B4 and GFP_P4 were computed using Equation 3 using reporters spliced from 10% to 80% efficiencies. Stabilities of full-length reporters were estimated using the regression model described in Equation 1. (E, F) The production enhancement effect is robust to alternative estimates of GFP_B4 and GFP_P4 spliced isoform stabilities. In (E), $\frac{\beta_s}{\alpha_0}$ values were estimated using reporters spliced between 10% to 70% efficiency. In (F), $\frac{\beta_s}{\alpha_0}$ values were estimated using reporters spliced between 10% to 90% efficiency.

### Figure S5

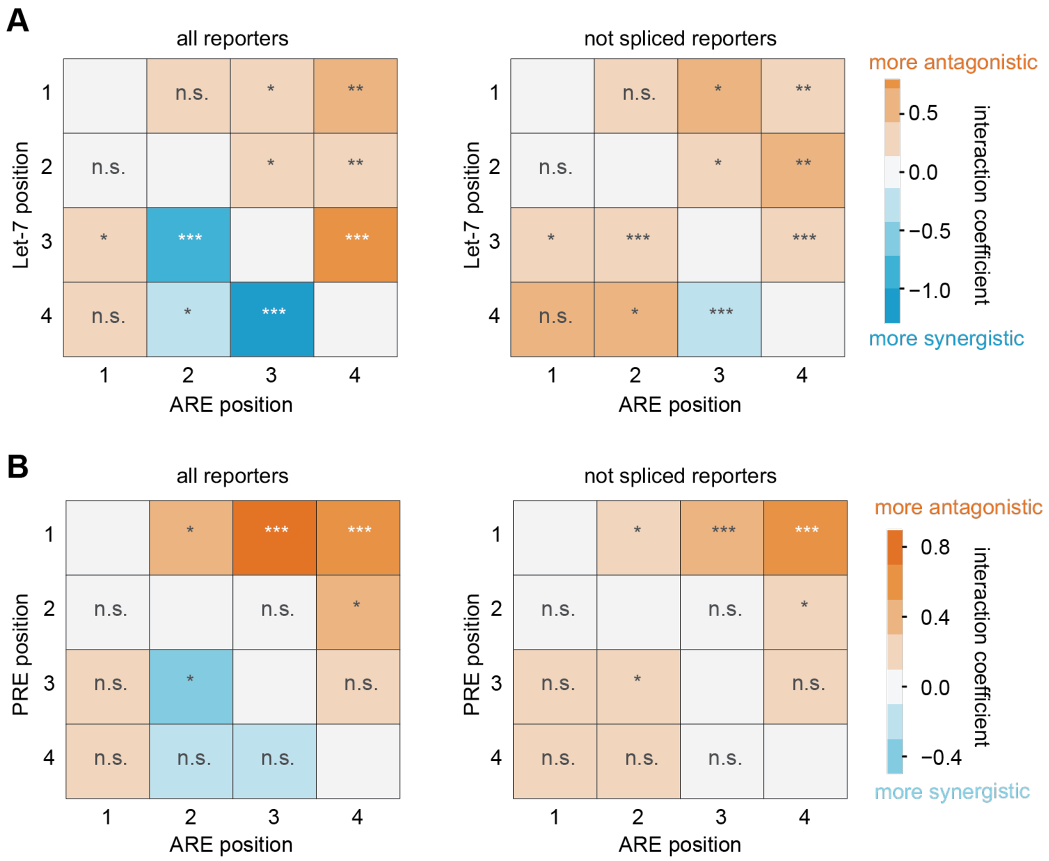


**Figure S5. Impact of AREs on neighboring Let-7 and PRE sites.** Interaction coefficients between (A) ARE and Let-7 modules and (B) ARE and PRE modules obtained from fitting a linear regression model (Equation 1) to all reporters or only to unspliced reporters (<1% spliced). In the original PTRE-seq publication (ref. 24), AREs were concluded to have both synergistic and antagonistic relationship with other regulatory modules depending on their positions. However, when spliced reporters are excluded, the regression model indicates that AREs have no impact or moderately antagonistic impact on repression by neighboring Let-7 and PRE sites. Statistical significance was assessed using the Wald test, where *=$P$ < 0.05, **=$P$ < 0.01, ***=$P$ < 0.001, and n.s.=not significant.

### Figure S6

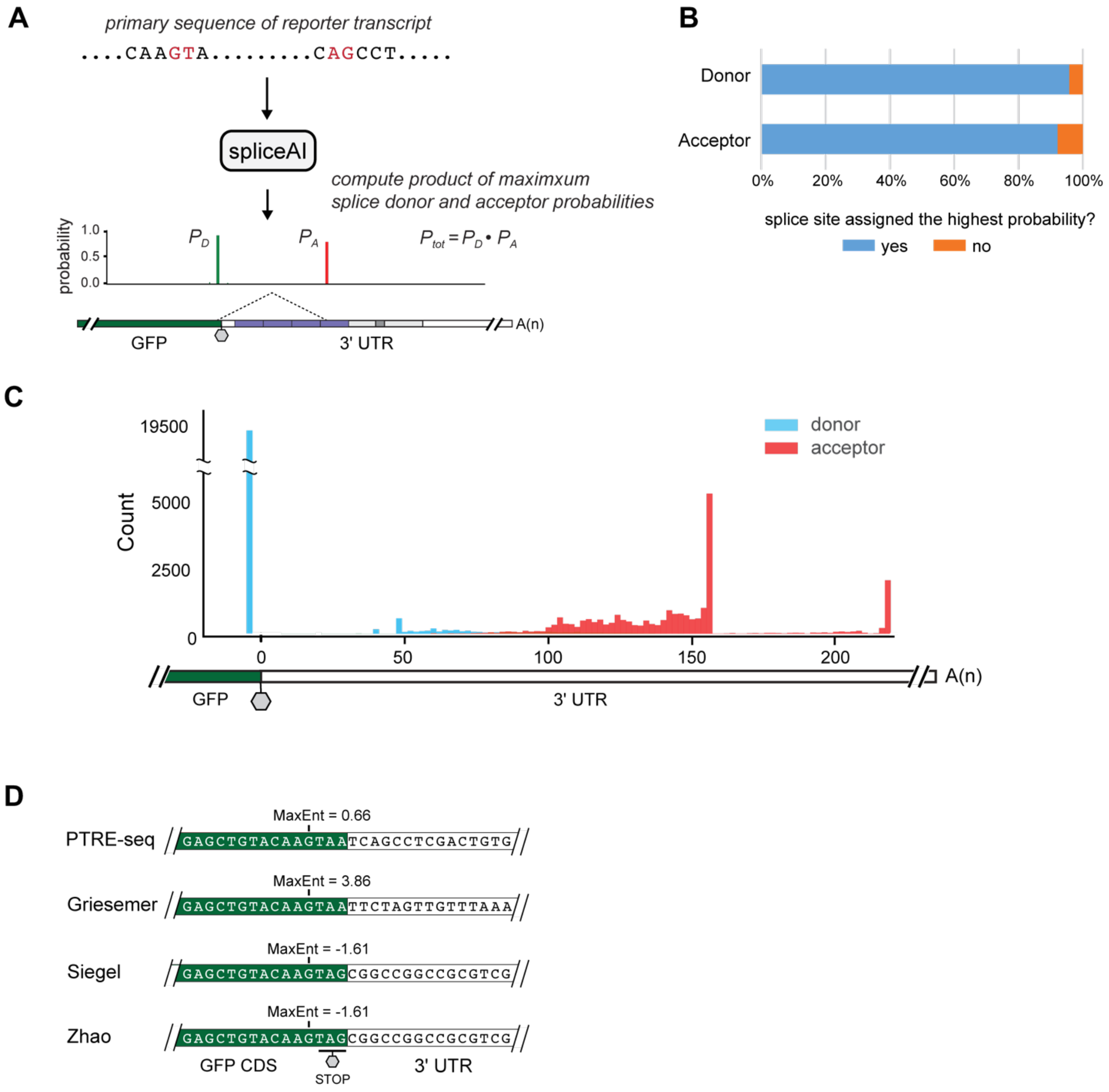


**Figure S6. Prediction of cryptic splicing in MPRAs using spliceAI.** (A) Strategy for predicting cryptic splicing in reporter transcripts using spliceAI. The transcript sequence of each reporter is input into spliceAI[36], which computes the probability of every nucleotide being a splice donor or acceptor. To compute the overall probability of splicing, we compute the product of the highest probability donors and acceptors, respectively. (B) Percentage of splice sites in PTRE-seq with usage >1% correctly assigned as the highest probability sites by spliceAI. (C) Relative location of predicted 5' and 3' splice sites in *Griesemer*[7], *Siegel*[8], and *Zhao*[9] MPRAs. Only splice sites with probability above 0.01 are included. (D) Sequence differences in the GFP cryptic donor site in the reporter vectors used for PTRE-seq, *Griesemer*, *Siegel*, and *Zhao*. GFP donor is scored using Maximum Entropy[52].

**Figure S7**

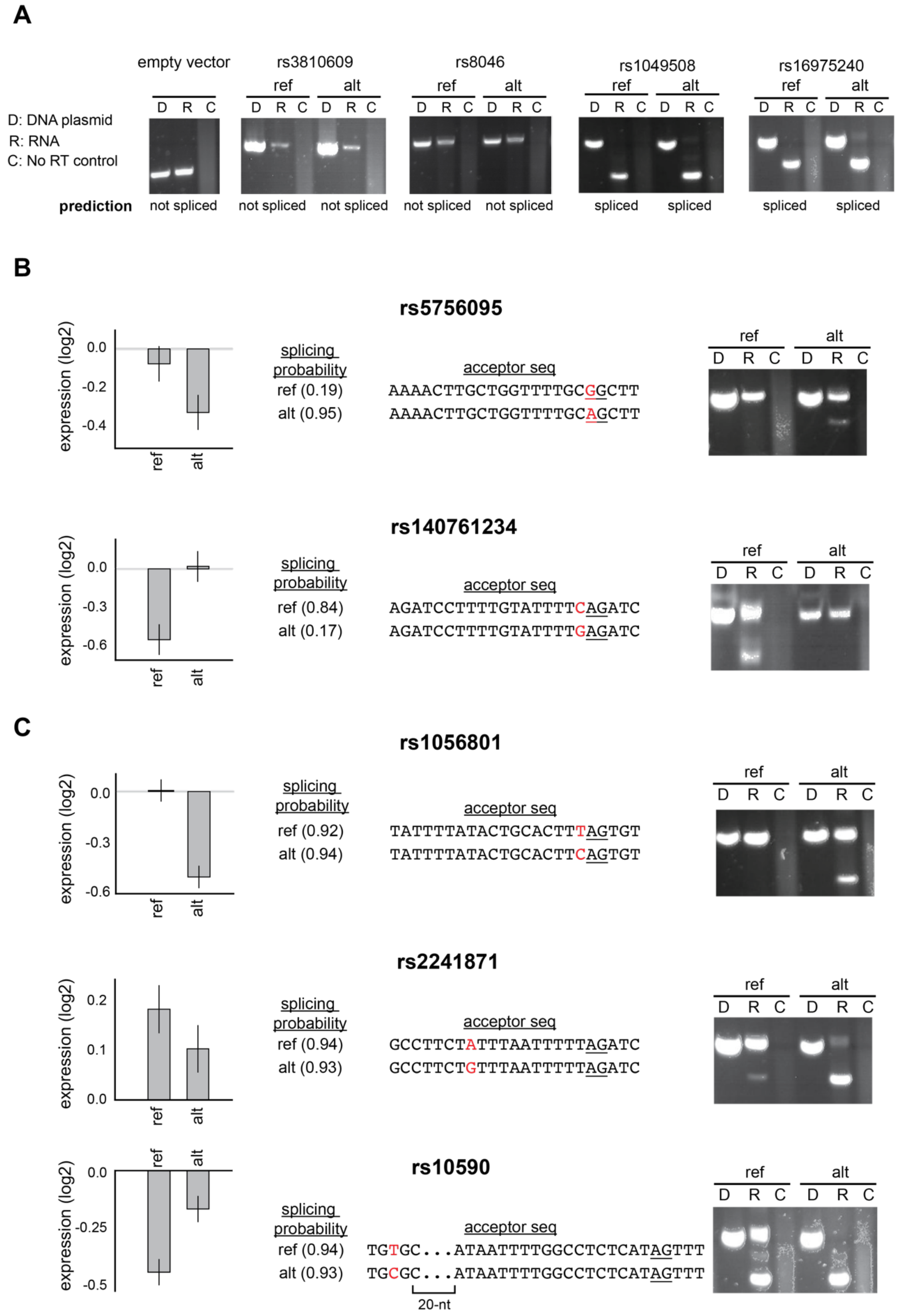
A
empty vector
rs3810609
rs8046
rs1049508
rs16975240
ref
alt
D R C
D: DNA plasmid
R: RNA
C: No RT control
prediction
not spliced
spliced
B
rs5756095
expression (log2)
0.0
-0.2
-0.4
splicing probability
ref (0.19)
alt (0.95)
acceptor seq
AAAACTTGCTGGTTTTGCGGCTT
AAAACTTGCTGGTTTTGCAGCTT
rs140761234
0.0
-0.3
-0.6
ref (0.84)
alt (0.17)
AGATCCTTTTGTATTTTCAGATC
AGATCCTTTTGTATTTTGAGATC
C
rs1056801
ref (0.92)
alt (0.94)
TATTTTATACTGCACTTTAGTGT
TATTTTATACTGCACTTCAGTGT
rs2241871
0.2
0.1
0.0
ref (0.94)
alt (0.93)
GCCTTCTATTTAATTTTTAGATC
GCCTTCTGTTTAATTTTTAGATC
rs10590
0.0
-0.25
-0.5
ref (0.94)
alt (0.93)
TGTGC...ATAATTTTGGCCTCTCATAGTTT
TGCGC...ATAATTTTGGCCTCTCATAGTTT
20-nt

**Figure S7. Validation of cryptic splicing in *Greisemer* MPRA.** Selected reporters from the *Griesemer* MPRA[7] were synthesized and validated with RT-PCR analysis. (A) Agarose gels of plasmid backbone, 2 tamVar pairs predicted to both be unspliced, and two tamVar pairs predicted to both be highly spliced. (B) Analysis of tamVar pairs predicted to be differentially spliced. See also Figure 5H. (C) Analysis of tamVar pairs in which both alleles were predicted to be strongly spliced, but RT-PCR indicated differential splicing. For (B) and (C), left bar plots show mean relative expression measured by *Griesemer*. Error bars denote standard error. SpliceAI probability for each reporter and sequence of predicted 3' splice sites are shown in the middle. Variant is highlighted red and the splice acceptor is underlined. At right is shown RT-PCR analysis of individually transfected plasmids resolved by agarose gel. D: DNA plasmid, R: RT-PCR of RNA, C: no RT control.

#### Table S1: Primers used in this study

| Name | Sequence |
|---|---|
| RE_Amp_RT | AAAGGACAGTGGGAGTGGC |
| GFP_Amp_F_seq | GACTGGAGTTCAGACGTGTGCTCTTCCGATCTNNNNNGCTGCCCGACAACCAC |
| RE_Amp_R_seq | CCCTACACGACGCTCTTCCGATCTNNNNNAAAGGACAGTGGGAGTGGC |
| Junction_R | CACAGTCGAGGTTGTACAGCTC |
| M1_For | GAGCTATACAAATAATCAGCCTCGACTGTGGC |
| M1_Rev | GCCACAGTCGAGGCTGATTATTTGTATAGCTC |
| M2_For | GAGCTATACCAAATAATCAGCCTCGACTGTGGC |
| M2_Rev | GCCACAGTCGCCGGCCGCTACTTGTATAGCTC |
| Griesemer_Amp_F | CTAGTTGTTTAAAGCCCAACGCTAG |
| Griesemer_Amp_R | CGACGACGCTCTTCCGATC |
| Griesemer_F | GTTTAAAGCCCAACGCTAGTC |
| Griesemer_RT | CAGCGGTGGCAGCAG |
| Griesemer_F_seq | GACTGGAGTTCAGACGTGTGCTCTTCCGATCTNNNNNGTTTAAAGCCCAACGCTAGTC |
| Griesemer_R_seq | CCCTACACGACGCTCTTCCGATCTNNNNNCAGCGGTGGCAGCAG |

#### Table S2: Evaluated sequences from the *Griesemer*[7] MPRA

Sequences were synthesized as IDT eBlocks, which have a minimum length requirement of 300 nts. To meet this minimum length requirement, the following sequence was appended 3' to all sequences during synthesis:

TCGGATCCCGAAGAACTGCGCAGTCCGGTTATAACGTCTGACATGTCCTTTTTTGATTTGGAATCCAACC
ACTCAAGTGGATCCGTTCATTTACTTTGCGAAAAATATACCCACAAATTGCCCATCGAAAGTGA

This flanking sequence was then removed by the initial PCR amplification step (Methods).

| Name | Sequence |
|---|---|
| rs8046_ref | CTAGTTGTTTAAAGCCCAACGCTAGTCACTTTCCGAGCTCGCTAGCCTCCAACAAGAATG CATTCCCTGAATCTGTGCCTGCACTGAGAGGGCAAGGAAGTGGGGTGTTCTTCTTGGGAC CCCCACTAAGACCCTGGTCTGAGGATGTAAGATCGGAAGAGCGTCG |
| rs8046_alt | CTAGTTGTTTAAAGCCCAACGCTAGTCGAAAGTCGAGCTCGCTAGCCTCCAACAAGAATG CATTCCCTGAATCTGTGCCTGCACTGAGAGGGCAAGGAGGTGGGGTGTTCTTCTTGGGAC CCCCACTAAGACCCTGGTCTGAGGATGTAAGATCGGAAGAGCGTCG |
| rs140761234_ref | CTAGTTGTTTAAAGCCCAACGCTAGTCATGCCGCGAGCTCGCTAGCCTCTCCAGCCTGGG AAAAGTTCTCCTTATTTGTTTAGATCCTTTTGTATTTTCAGATCTCCTTGGAGCAGTAGA GTACCTGGTAGACCATAATAGTGGAAAAGAGATCGGAAGAGCGTCG |
| rs140761234_alt | CTAGTTGTTTAAAGCCCAACGCTAGTCCGGCATCGAGCTCGCTAGCCTCTCCAGCCTGGG AAAAGTTCTCCTTATTTGTTTAGATCCTTTTGTATTTTGAGATCTCCTTGGAGCAGTAGA GTACCTGGTAGACCATAATAGTGGAAAAGAGATCGGAAGAGCGTCG |
| rs5756095_ref | CTAGTTGTTTAAAGCCCAACGCTAGTCCGGCTACGAGCTCGCTAGCCTTACACTCCCTCC CCTTTTGAAAGTCCCTAATAAAAACTTGCTGGTTTTGCGGCTTGTGAGGCATCACGGAAC CTACCGATGTGTGATGTCTCCCCTGGACAAGATCGGAAGAGCGTCG |

| | |
|---|---|
| rs5756095_alt | CTAGTTGTTTAAAGCCCAACGCTAGTCTAGCCGCGAGCTCGCTAGCCTTACACTCCCTCC<br>CCTTTTGAAAGTCCCTAATAAAAACTTGCTGGTTTTGCAGCTTGTGAGGCATCACGGAAC<br>CTACCGATGTGTGATGTCTCCCCTGGACAAGATCGGAAGAGCGTCG |
| rs1056801_ref | CTAGTTGTTTAAAGCCCAACGCTAGTCCGTACTCGAGCTCGCTAGCCTAGCAGTTGGTCT<br>ATTCAGAATCAAACCTTTTTATATTTTATACTGCACTTTAGTGTATTTTCTGTCACTGTA<br>GGTATAGAAGATCTGCCTCCCCTGTGGAAAGATCGGAAGAGCGTCG |
| rs1056801_alt | CTAGTTGTTTAAAGCCCAACGCTAGTCAGTACGCGAGCTCGCTAGCCTAGCAGTTGGTCT<br>ATTCAGAATCAAACCTTTTTATATTTTATACTGCACTTCAGTGTATTTTCTGTCACTGTA<br>GGTATAGAAGATCTGCCTCCCCTGTGGAAAGATCGGAAGAGCGTCG |
| rs1049508_ref | CTAGTTGTTTAAAGCCCAACGCTAGTCCGTGTCCGAGCTCGCTAGCCTTTTTCATATACA<br>TCTTACCTCATTTCAAGTGAATTATTTTAATCTTTTTCTCTCTTTCCAAAAATTTACAGG<br>AATGTTTAGTGTAATTGGATTTCGCTATCAGATCGGAAGAGCGTCG |
| rs1049508_alt | CTAGTTGTTTAAAGCCCAACGCTAGTCGACACGCGAGCTCGCTAGCCTTTTTCATATACA<br>TCTTACCTCATTTCAAGTGAATTATTTTAATCTTTTCCTCTCTTTCCAAAAATTTACAGG<br>AATGTTTAGTGTAATTGGATTTCGCTATCAGATCGGAAGAGCGTCG |
| rs2241871_ref | CTAGTTGTTTAAAGCCCAACGCTAGTCCTTCTTCGAGCTCGCTAGCCTTCTTGCCATAAA<br>TTGCTGACTTCACGGTTAGGTAAAGATGGATAGATAAAGAGAAGCTTAAACAGTGGCATT<br>TTACTACAAAGGCCTTCTATTTAATTTTTAGATCGGAAGAGCGTCG |
| rs2241871_alt | CTAGTTGTTTAAAGCCCAACGCTAGTCAAGAAGCGAGCTCGCTAGCCTTCTTGCCATAAA<br>TTGCTGACTTCACGGTTAGGTAAAGATGGATAGATAAAGAGAAGCTTAAACAGTGGCATT<br>TTACTACAAAGGCCTTCTGTTTAATTTTTAGATCGGAAGAGCGTCG |
| rs13004845_ref | CTAGTTGTTTAAAGCCCAACGCTAGTCATGAACCGAGCTCGCTAGCCTTTTTTTTTTTTC<br>TTTACATTGGCTTTTTAAAAGACAGTTTTTATTTTTCAGATACTGGAATGTCAAGTATCT<br>GGGTTGTCCCAAGACGTTGGAGATAATTCAGATCGGAAGAGCGTCG |
| rs13004845_alt | CTAGTTGTTTAAAGCCCAACGCTAGTCGTTCATCGAGCTCGCTAGCCTTTTTTTTTTTTC<br>TTTACATTGGCTTTTTAAAAGACAGTTTTTATTTTTCAAATACTGGAATGTCAAGTATCT<br>GGGTTGTCCCAAGACGTTGGAGATAATTCAGATCGGAAGAGCGTCG |
| rs3810609_ref | CTAGTTGTTTAAAGCCCAACGCTAGTCAATCACCGAGCTCGCTAGCCTCGTCTCAGCATC<br>GCCAGTCTAGACTGTCTATGGAGAGCAGAAAGTTGTCTGGGGCTGCCTGGGGAACTGTGA<br>GGCCAGCTATATCACCGTCGCTGATGGTGAGATCGGAAGAGCGTCG |
| rs3810609_alt | CTAGTTGTTTAAAGCCCAACGCTAGTCGTGATTCGAGCTCGCTAGCCTCGTCTCAGCATC<br>GCCAGTCTAGACTGTCTATGGAGAGCAGAAAGTTGTCTAGGGCTGCCTGGGGAACTGTGA<br>GGCCAGCTATATCACCGTCGCTGATGGTGAGATCGGAAGAGCGTCG |
| rs16975240_ref | CTAGTTGTTTAAAGCCCAACGCTAGTCGCAGGTCGAGCTCGCTAGCCTCGACGTAGGGTA<br>AGTGCAATCCCAAGCCGTTTAAAATAATCCCAGACTGCCTGGAGGCTTTGTTCTTATTTT<br>CTGATTCTTTTTTCTTTGTCTTTGTTGGAAGATCGGAAGAGCGTCG |
| rs16975240_alt | CTAGTTGTTTAAAGCCCAACGCTAGTCACCTGCCGAGCTCGCTAGCCTCGACGTAGGGTA<br>AGCGCAATCCCAAGCCGTTTAAAATAATCCCAGACTGCCTGGAGGCTTTGTTCTTATTTT<br>CTGATTCTTTTTTCTTTGTCTTTGTTGGAAGATCGGAAGAGCGTCG |
| rs10590_ref | CTAGTTGTTTAAAGCCCAACGCTAGTCCTCTACCGAGCTCGCTAGCCTTTGTGTATGCTT<br>CATCACCTATATTAGGCAAATTCCATTTTTTTCCCTTGTGCTAAGGTAAAGATTTAATTA<br>AATAATTTTGGCCTCTCATAGTTTTCTCTAGATCGGAAGAGCGTCG |
| rs10590_alt | CTAGTTGTTTAAAGCCCAACGCTAGTCGTAGAGCGAGCTCGCTAGCCTTTGTGTATGCTT<br>CATCACCTATATTAGGCAAATTCCATTTTTTTCCCTTGCGCTAAGGTAAAGATTTAATTA<br>AATAATTTTGGCCTCTCATAGTTTTCTCTAGATCGGAAGAGCGTCG |
